# Optical segmentation-based compressed readout of neuronal voltage dynamics

**DOI:** 10.1101/2023.11.10.566599

**Authors:** Seonghoon Kim, Gwanho Ko, Iksung Kang, He Tian, Linlin Z. Fan, Yixin Li, Adam E. Cohen, Jiamin Wu, Qionghai Dai, Myunghwan Choi

## Abstract

Functional imaging of biological dynamics generally begins with acquiring time-series images, followed by quantifying spatially averaged intensity traces for the regions of interest (ROIs). The conventional pipeline discards a substantial portion of the acquired data when quantifying intensity traces, indicative of inefficient data acquisition. Here we propose a conceptually novel acquisition pipeline that assigns each ROI to a single pixel in the detector, enabling optimally compressed acquisition of the intensity traces. As a proof-of-principle, we implemented a detection module composed of a pair of spatial light modulators and a microlens array, which segments the original image into multiple subimages by introducing distinct angular shifts to each ROI. Each subimage exclusively encodes the signal for the corresponding ROI, facilitating the compressed readout of its intensity trace using a single pixel. This spatial compression allowed for maximizing the temporal information without compromising the spatial information on ROIs. Harnessing our novel approach, we demonstrate the recording of circuit-scale neuronal voltage dynamics at over 5 kHz sampling rate, revealing the individual action potential waveforms within subcellular structures, as well as their submillisecond-scale temporal delays.

## Introduction

Temporal dynamics of neuronal membrane potentials serve as the fundamental signal in neural computation^1^. In particular, action potentials encode rich information in their millisecond-scale analogue waveforms, such as the composition of ion channels, electrical excitability, and neural connectivity, which are indispensable for understanding neuronal physiology and pathology^2–5^. Patch clamp recording, which measures the electrical properties of individual neurons with an electrode, has long been the gold-standard tool for studying neuronal waveforms due to its unparalleled temporal resolution and signal-to-noise ratio (SNR). However, its experimental throughput is low, typically limited to one neuron at a time, and targeting the fine processes of neurons is often infeasible due to the physical size of the electrode^6^. These limitations hinder a deeper understanding of neuronal dynamics at a circuit level.

With recent advances in fluorescent voltage indicators, voltage imaging has gained interest since it provides a non-contact optical readout on voltage dynamics with subcellular-scale spatial resolution^7–15^. This unique capability, in principle, allows for kilohertz-scale recording of voltage dynamics in neural processes, as well as cell bodies, in a large neural population^9,10,16–21^. However, simultaneously achieving the subcellular-level spatial and submillisecond-scale temporal resolutions over the wide field-of-view remains challenging due to the technical limitations of optical detectors^22^. For example, increasing a camera’s acquisition speed is achieved by compromising spatial information, either by decreasing the field-of-view (subarray readout) or reducing the spatial sampling rate (pixel binning). Consequently, achieving kilohertz-scale acquisition speed inevitably accompanies spatial crosstalk among adjacent objects or sacrifices field-of-view, which poses a fundamental challenge in capturing complex dynamics from spatially entangled neural circuits.

For measuring functional neural dynamics, time-series images are generally acquired first and the intensity traces from the selected regions of interest (ROIs) are subsequently analyzed by spatial averaging (**Fig. 1a**)^20,23–27^. In this conventional pipeline, the data size of the acquired time-series images is proportional to the number of pixels in the image multiplied by the number of frames. For instance, an acquisition of 20 neurons with 512 × 512 pixels and a 1 kHz frame rate at a 16-bit depth exceeds 30 GB in a minute. In contrast, the data size of the resulting intensity traces for the 20 neurons is merely ∼2.4 MB, corresponding to less than 0.01% of the acquired data. In this regard, the conventional pipeline is deemed inefficient, because a large portion of the acquired data, obtained at the cost of compromised temporal resolution, is discarded during the analysis^28,29^.

**Fig. 1:**
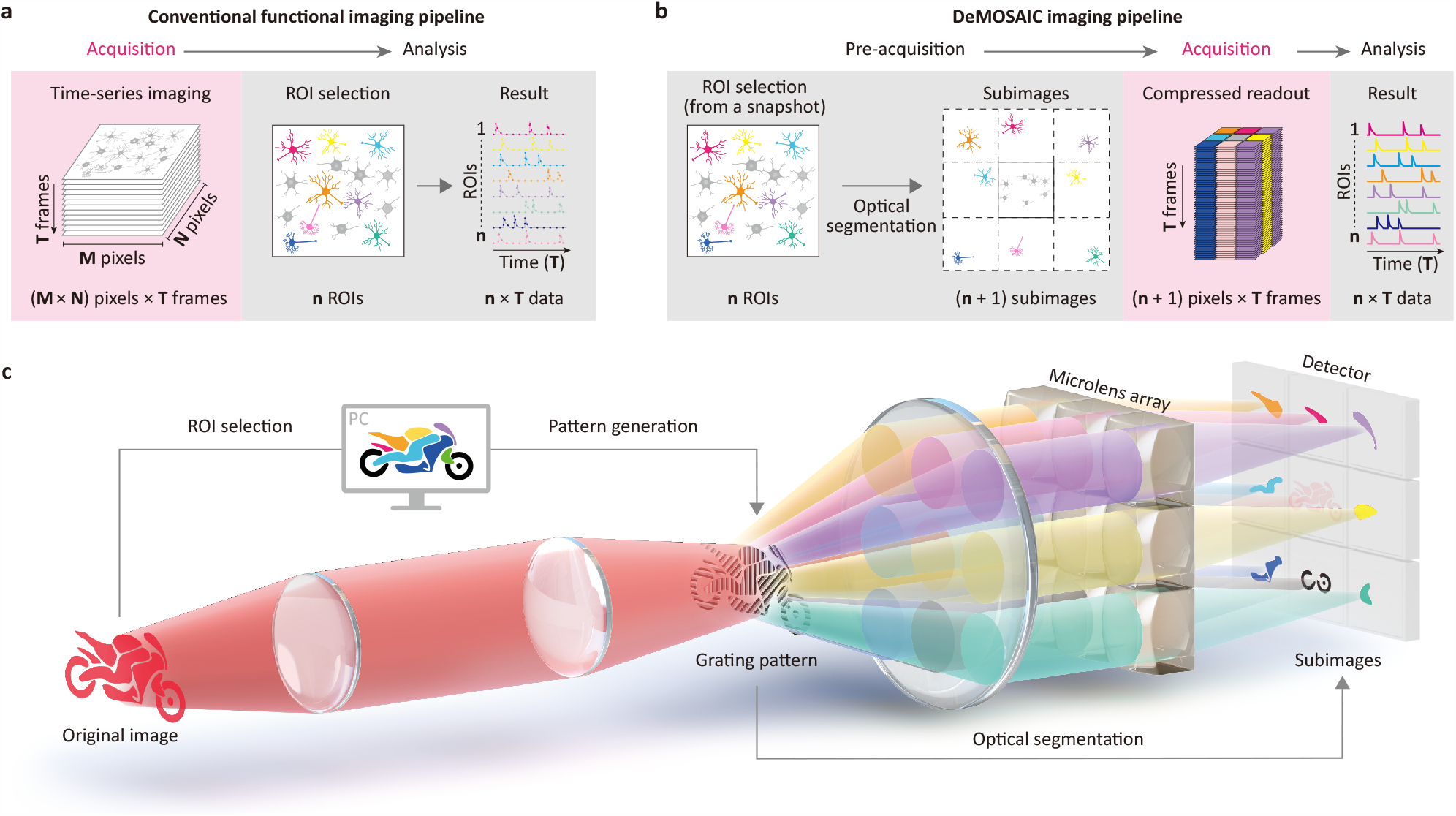
Principle of DeMOSAIC. **a**, A conventional functional imaging pipeline. Time-series images are acquired first (image block size: M × N × T), and then intensity traces for the selected n ROIs are obtained, resulting in ‘n × T’ data. **b**, The proposed DeMOSAIC imaging pipeline. The ROIs are selected from a digital snapshot image prior to the acquisition. The subimages corresponding to individual ROIs are optically segmented from the original image, and are assigned to each pixel in a detector. Thus, the dimension of the acquired data closely matches with that of the resulting data. **c**, A conceptual illustration of the DeMOSAIC acquisition. The original image is optically relayed to the ROI-based blazed grating pattern, which provides distinct angular modulations for each ROI. Subsequently, the second optical relay including a microlens array (MLA) projects each ROI-based subimage to each pixel in a detector.

To address the limitation of the conventional pipeline, we developed a conceptually novel detection scheme named DeMOSAIC (Diffractive Multisite Optical Segmentation Assisted Image Compression), in which only a single pixel is assigned for each ROI so that the data dimension of the acquisition matches that of the resulting data (**Fig. 1b**). In the DeMOSAIC pipeline, the ROIs are preselected from a snapshot image prior to time-series acquisition, and the field-of-view is optically divided into a number of subimages configured to contain individual ROIs. Consequently, the number of pixels closely matches the number of ROIs, providing optimal efficiency in data size. We first demonstrate the implementation of the DeMOSAIC system providing optical segmentation by introducing a patterned blazed grating and a microlens array along the detection path (**Fig. 1c**). Second, we show a proof-of-principle that DeMOSAIC effectively realizes compressive recording at >100 kHz without introducing spatial crosstalk. Lastly, we demonstrate subcellular-scale voltage imaging of neural circuit dynamics at a sampling rate of >5 kHz.

## Results

### Implementation of the DeMOSAIC system

We implemented the DeMOSAIC system as an add-on to the detector port of an inverted epifluorescence microscope (**Fig. 2a, Extended Data Fig. 1** and **Supplementary Table 1**). The sCMOS camera mounted in the original detection path provided a snapshot image for the userdefined selection of ROIs. Following ROI selection, we redirected the detection beam path to the DeMOSAIC system using a motorized flip mirror, which relayed the original image to the spatial light modulators (SLM) for optical segmentation. In the meanwhile, we generated the grating pattern for the SLMs by assigning one of eight 3-level blazed grating patterns to each ROI, providing directional first-order diffraction at an angle of ∼1° (**Fig. 2a** and **Extended Data Fig. 2**).

**Figure 2:**
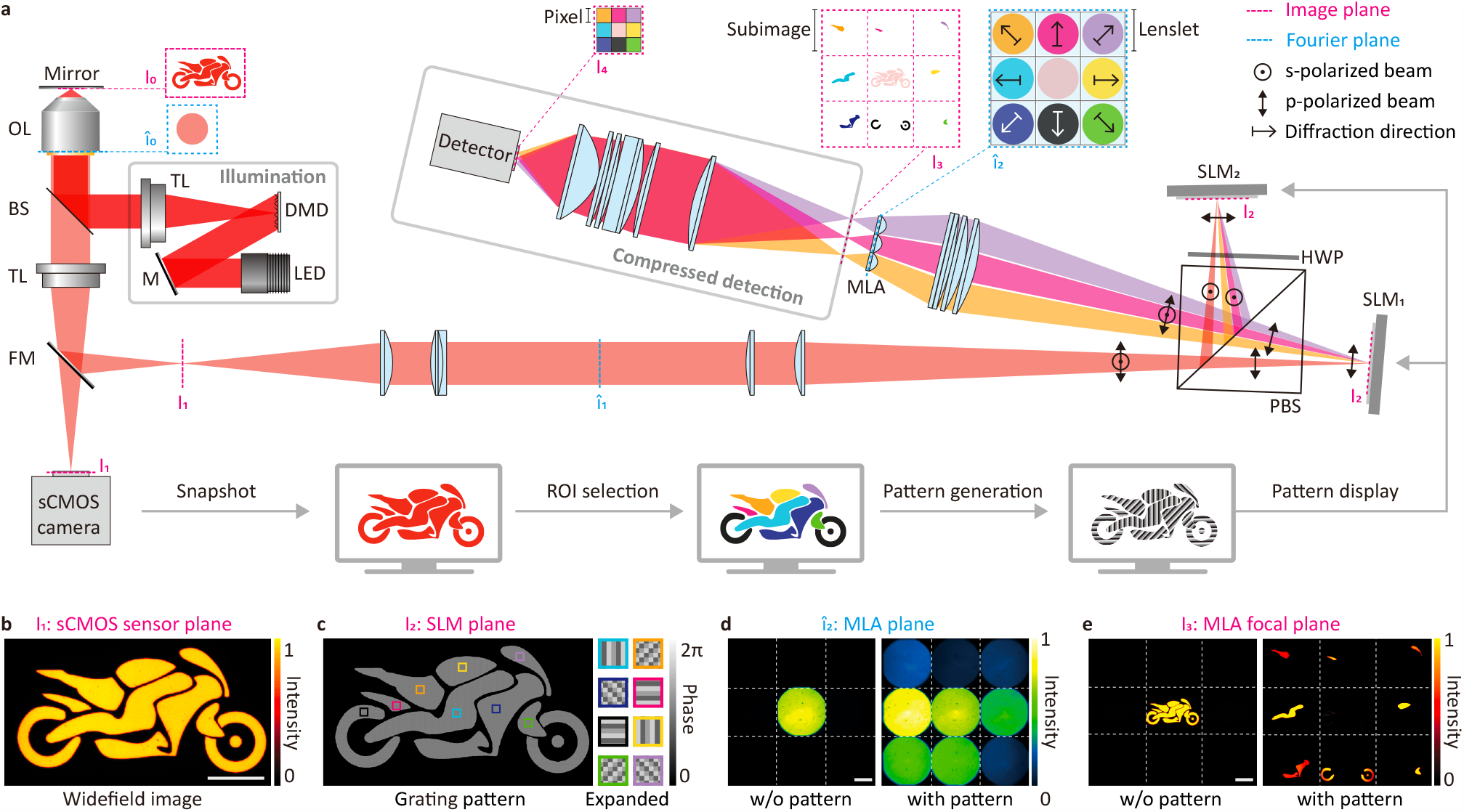
Instrumentation of the DeMOSAIC system. **a**, Schematic optical layout of the DeMOSAIC system. A motorcycle-shaped light pattern was generated by a digital micromirror device (DMD) coupled with an LED and projected onto the mirror at the sample plane (I_0_). The widefield image is captured by an sCMOS camera for ROI selection (I_1_) and is subsequently relayed to the SLMs using a motorized flip mirror (FM). Using a pair of SLMs at the conjugate image plane (I_2_), original image is optically segmented by introducing a distinct angular shift. The diffracted beams are refocused by a microlens array (MLA) to form 3-by-3 subimages (I_3_), each of which encodes the individual ROI. The subimage plane is optically demagnified and relayed to a detector (I_4_) in the compressed detection module. OL, objective lens. TL, tube lens. BS, beam splitter. LED, light emitting diode. M: mirror. PBS, polarizing beam splitter; HWP, half-waveplate. **b**, A reflectance widefield image of the motorcycle-shaped pattern. Scalebar, 50 μm. **c**, A grating pattern generated based on the selected ROIs. The 8 types of pattern units are used. **d**, Image of the Fourier plane at MLA plane (î_2_), with and without pattern displayed on SLM as indicated. Scalebar, 1 mm. **e**, The subimage plane (I_3_) with and without pattern displayed on SLM. Scalebar, 1 mm.

To impose phase modulation to unpolarized fluorescence emission, we divided the emission signal into s- and p-polarized beams by a polarizing beam splitter (PBS) and introduced a pair of polarization-sensitive reflective SLMs for each polarization (**Fig. 2a** and **Extended Data Fig. 1**). In the beam path for the s-polarized beam, we placed a half-wave plate at 45° to rotate the polarization by 90°. To account for the reflection geometry at the PBS, we configured the input grating pattern for the s-polarized beam to be inverted with respect to that for the p-polarized beam. We coregistered the image planes of the SLMs and the original image plane using the space transformation matrices obtained by the point-based multimodal registration algorithm (**Extended Data Fig. 3**). Consequently, we achieved distinct angular shifts for up to 8 individual ROIs, while the remaining non-selected background remained in the zeroth order.

After angular modulation of the ROIs, we introduced another image relay system comprising a microlens array (MLA) to segregate individual ROIs into subimages. The MLA was fabricated by lasercutting of nine plano-convex lenses into 2.8-by-2.8 mm^2^ square-shaped lenslets and assembling them into a 3-by-3 square grid using optical adhesive (**Extended Data Fig. 4**). This image relay system provided the subimage plane (I_3_), composed of the central background subimage and the surrounding eight ROI-based subimages. Each subimage exclusively encodes the signal of the selected ROI, thus the intensity trace for each ROI can be compressively recorded by a single pixel.

Having implemented the DeMOSAIC system, we evaluated its feasibility for optical segmentation using a synthetic sample image. Using a digital micromirror device (DMD) coupled to a red light-emitting diode (LED), we projected a motorcycle-shaped light pattern onto a mirror surface positioned at the sample plane. We captured a widefield image with an sCMOS camera (**Fig. 2b**) and generated the ROI-based grating pattern to segment the motorcycle into eight distinct parts (**Fig. 2c**). Next, the flip mirror redirected the detection path to the DeMOSAIC beam path. Without the grating pattern, all the signal was in the zeroth order (**Fig. 2d,e**). After displaying the ROI-based grating pattern on the SLMs, the zeroth-order signal was segmented and redistributed towards the selected first-orders with high precision (**Fig. 2d,e**, **Extended Data Fig. 5**, and **Supplementary Section 1**). Other than the user-defined selection of ROIs, the process of DeMOSAIC acquisition is automated and is completed within several seconds.

### Demonstration of DeMOSAIC acquisition at 125 kHz

Given that DeMOSAIC acquisition provides optical segmentation enabling compressed readout, we designed an experiment to demonstrate its high-speed capability (**Fig. 3a**). First, we created a set of graffiti letters (D, E, M, O, S, A, i, C) and loaded them onto the DMD memory in the sequence of ‘i AM CODES’ and displayed each letter sequentially at the maximum refresh rate of 9.5 kHz (**Fig. 3b**). The sequential light patterns were projected onto a mirror surface, and the reflected signal was recorded via DeMOSAIC acquisition. This synthetic sample poses a challenge for conventional imaging approaches since it requires for both high spatial resolution to distinguish the intertwined graffiti letters and high temporal resolutions exceeding >19 kHz to avoid temporal aliasing.

**Figure 3:**
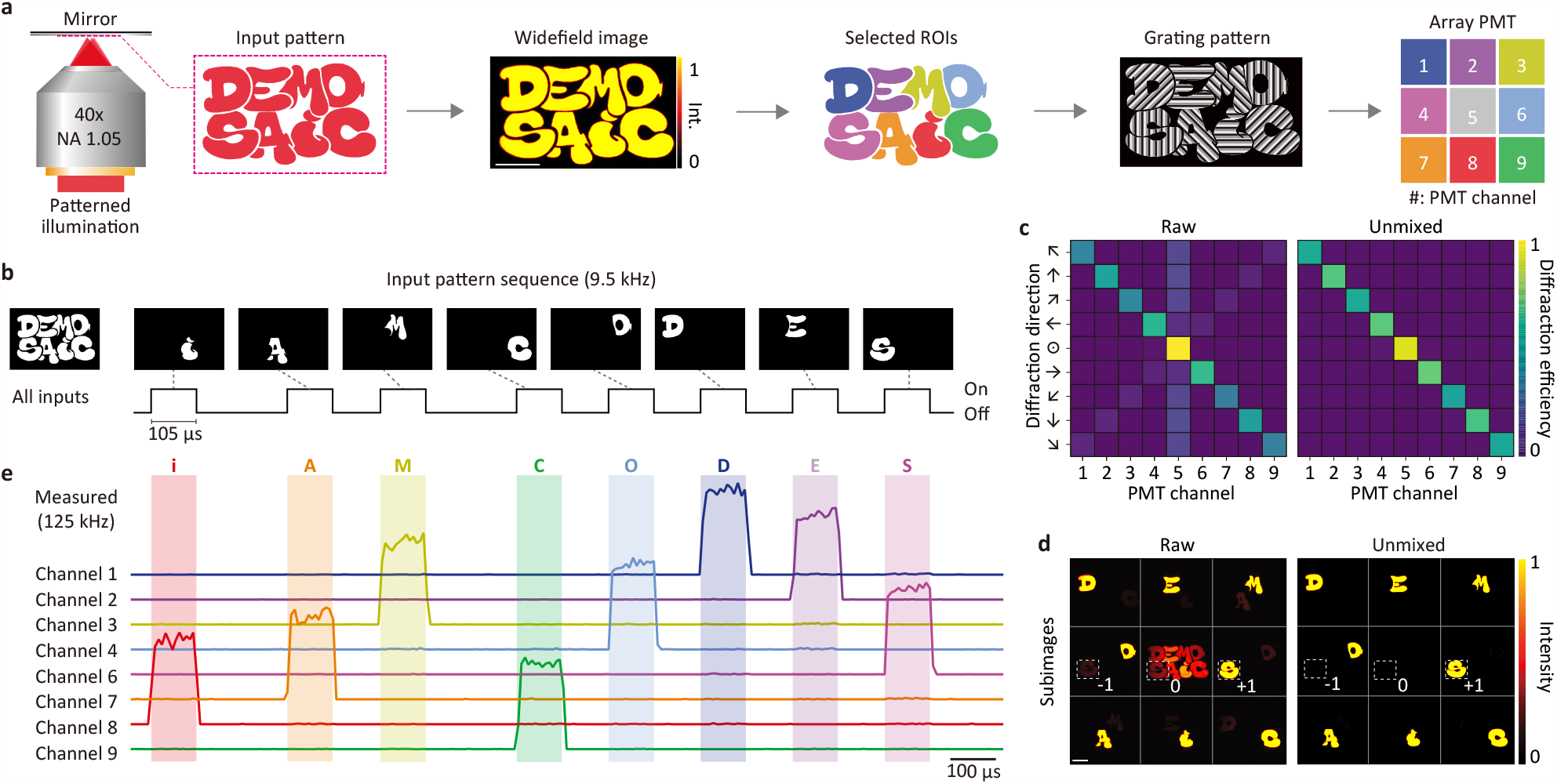
Ultrafast acquisition of a dynamic image via DeMOSAIC. **a**, The experimental procedures of the DeMOSAIC acquisition equipped with a PMT array detector. Each letter in the graffiti image was selected, segmented, and assigned to each PMT channel. Scalebar, 200 μm. **b**, Dynamic graffiti image. Using the binary pattern display mode of the DMD, each letter is displayed sequentially at a refresh rate of 9.5 kHz (105 μs for each letter with 105 or 210 μs intervals). **c**-**d**, Linear unmixing of interchannel crosstalk. Inter-channel crosstalk by −1^st^ order signals is evident in the raw diffraction efficiency matrix and subimages (left). Linear unmixing algorithm relocates the 0^th^ and −1^st^ order signals to the desired 1^st^ order (right). The dashed boxes indicate the −1st, 0th, and +1st orders for the letter ‘S’, as indicated. Scalebar, 200 μm. **e**, The 125-kHz readout of the dynamic graffiti pattern. The data was acquired by a 9-channel PMT array at 125 kHz and post-processed by applying the linear unmixing. The sequence is decoded as ‘i AM CODES’.

To capture the dynamic signal with a refresh rate of 9.5 kHz, we employed a PMT array connected to a 125-kHz digitizer as a detection module (**Extended Data Figs. 1** and **6**). After optical segmentation, however, we observed inter-channel crosstalk, which primarily arose from the −1st order diffractions (**Fig. 3 c,d**). The measured diffraction efficiencies for the 8 directions were 47.4 ± 11.9% for the +1st order, 13.6 ± 3.5% for the 0th order, 5.3 ± 0.3% for the −1st order, and the residual 32.8 ± 9.4%% for higher orders. Diffraction towards diagonal directions exhibited lower diffraction efficiency conceivably due to the pixelation of the grating. To resolve this issue, we employed a linear unmixing algorithm, which computationally redistributed the zeroth and −1st order signals to their corresponding +1st order channels (**Extended Data Fig. 7** and **Supplementary Section 2**). After applying the linear unmixing, the signal improved to 63.2 ± 8.8%, while diffraction crosstalk decreased from 5.3% to 0.3%. Consequently, we faithfully decoded the sequence of letters ‘i AM CODES’ by the DeMOSAIC acquisition with a temporal resolution of 8 µs (**Fig. 3e**).

### DeMOSAIC acquisition on neuronal calcium dynamics

We proceeded to apply the DeMOSAIC acquisition for imaging calcium dynamics of live neurons. To accomplish this, we adopted an EMCCD camera due to its high quantum efficiency, minimal read noise, and notably, the capability of analogue pixel binning. This capability, which is not supported by the alternative sCMOS camera, enabled us to customize pixel dimensions for optimized readout while preserving low read noise (**Extended Data Fig. 6**).

To capture functional dynamics over a large number of neurons, we optimized the acquisition pipeline (**Fig. 4**). First, we introduced pixel binning along the vertical axis using the ‘asymmetric binning mode’. In our demonstration, we used a subarray readout of 120 × 120 pixels and introduced a vertical pixel binning of 40 pixels. The overall pixel dimension was reduced to 120 × 3 pixels, with each subimage containing 40 × 1 pixels. This barcode-like subimage allowed to accommodate multiple ROIs in a subimage, particularly if the ROIs were sparsely distributed along the horizontal axis in a subimage. This configuration facilitated the simultaneous recording of over 20 neurons with DeMOSAIC acquisition. Second, we observed that the residual zeroth-order signal often saturated the sensor and interfered with the nearby subimages. To address this issue, we introduced patterned excitation, which selectively targets the excitation light to the signal-producing ROIs (**Extended Data Fig. 8**). The patterned excitation also offered advantages in recovering the zeroth-order signal through the linear-unmixing algorithm and minimizing photodamage to the neurons. Moreover, it enabled robust delineation of the optical segmentation boundary using the watershed algorithm (**Extended Data Fig. 5**).

**Figure 4:**
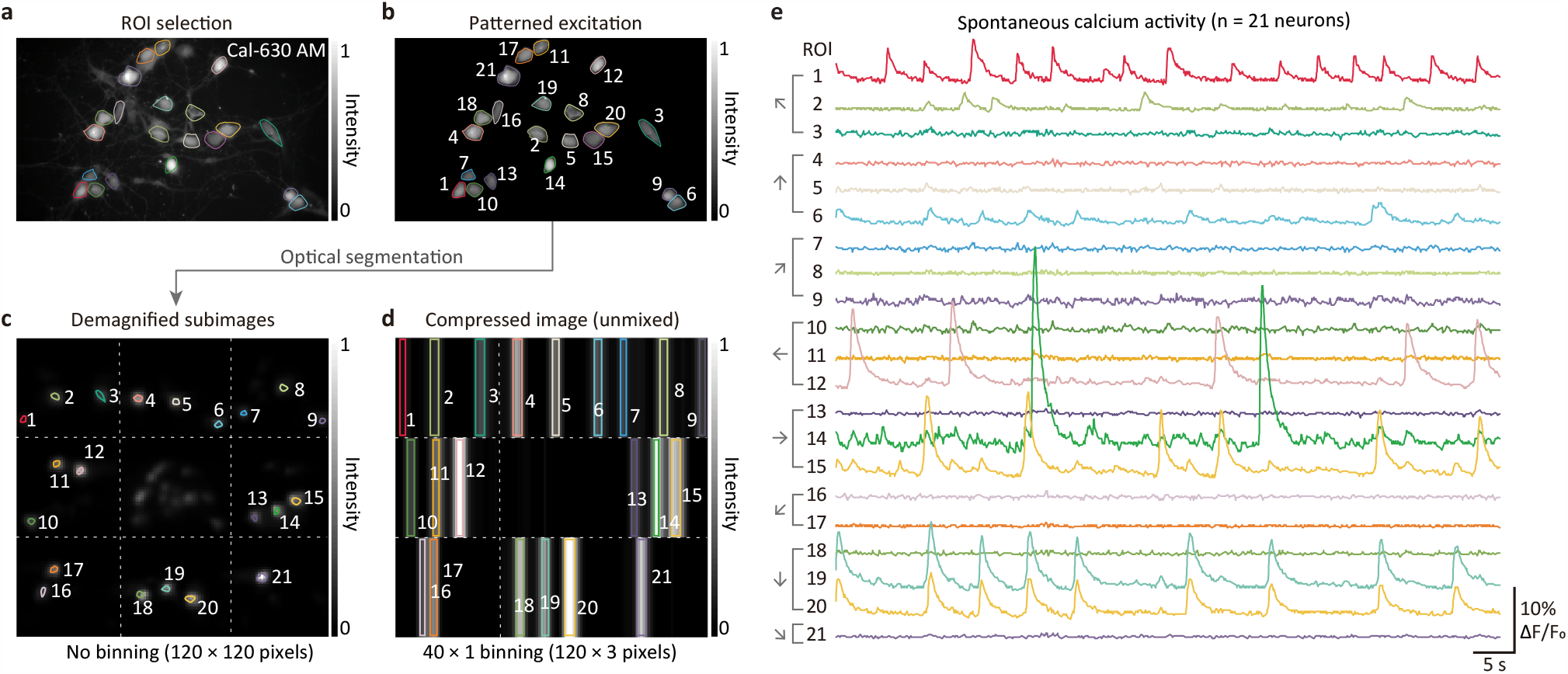
DeMOSAIC acquisition on neuronal calcium dynamics. **a**, A widefield fluorescence image of cultured neurons loaded with a calcium dye, Cal-630 AM. Scalebar, 50 μm. **b**, A fluorescence image with patterned excitation on the selected ROIs (n = 21 neurons). **c**, Segmented subimages taken by the EMCCD camera (120 × 120 pixels, 16 µm pixel size). **d**, Segmented subimages after vertical pixel binning by 40 pixels. The pixel dimension is compressed to 120 × 3 pixels, resulting in the barcode-like image. The image is displayed after applying the linear unmixing algorithm. **e**, Spontaneous neuronal calcium activity recorded by the DeMOSAIC pipeline (n = 21 neurons).

### DeMOSAIC acquisition on neuronal voltage dynamics

Harnessing the optimized DeMOSAIC acquisition pipeline, we observed subcellular-scale voltage dynamics in an intact neural circuit. We stained cultured neurons with a voltage-sensitive dye, BeRST1, providing excellent response kinetics and linearity to membrane potentials^14,15^. In a widefield fluorescence image, we selected 23 ROIs from three neurons, encompassing their neuronal processes and cell bodies. While recording their functional dynamics at a frame rate of 5.5 kHz, we periodically applied electric field stimulations (pulse width: 1 ms, pulse interval: 200 ms), sufficient to elicit action potentials in most neurons. Following linear unmixing, we extracted the voltage dynamics for each ROI and represented them as ‘dF/F_0_’ (**Fig. 5a**).

**Fig. 5:**
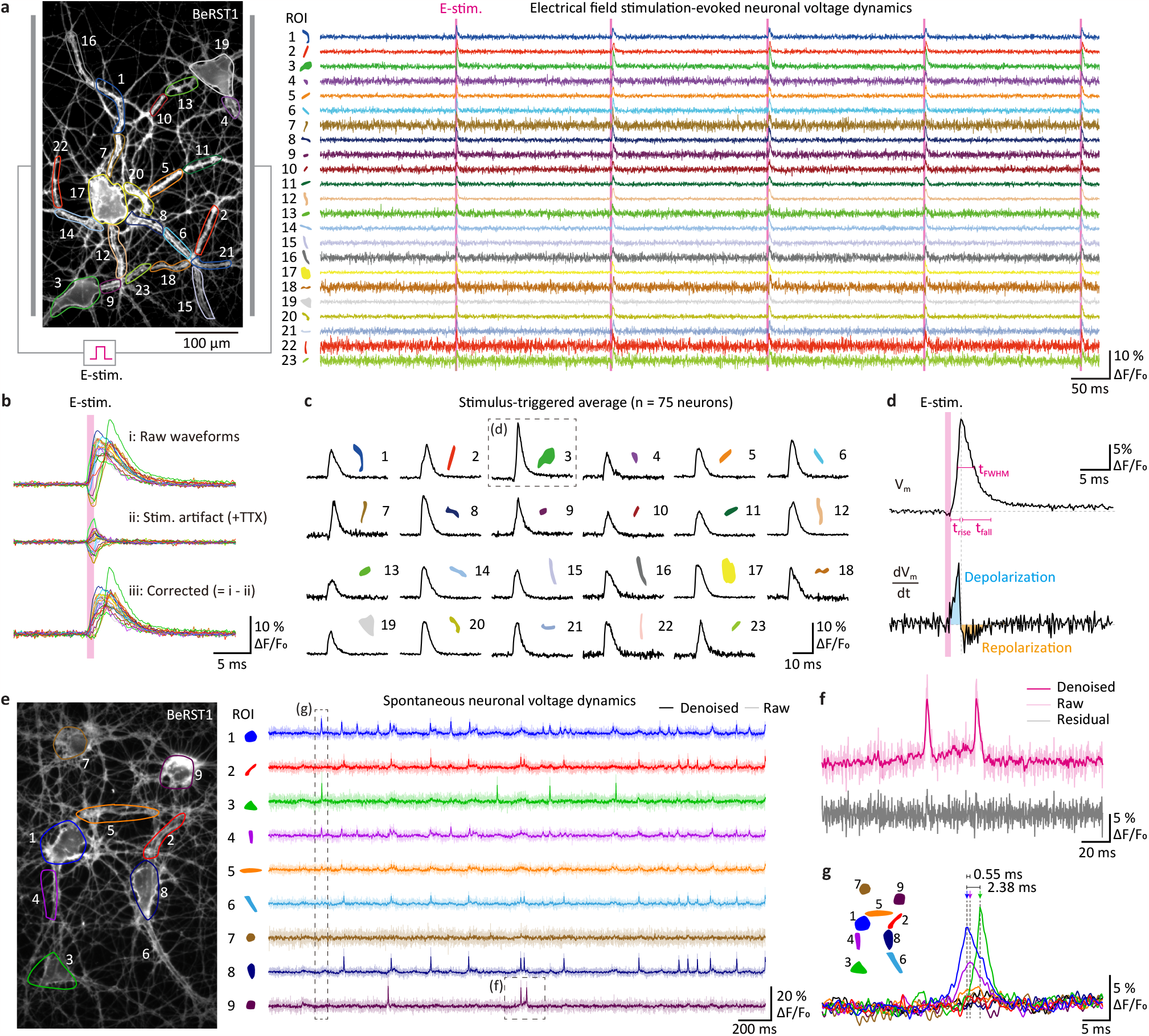
Recording subcellular-scale neuronal voltage dynamics at 5.5 kHz. **a**, Electrical stimulation evoked voltage dynamics recorded from the 23 ROIs at a frame rate of 5.5 kHz. The shaded areas indicate the electric field stimulations. **b**, Stimulus-triggered averaging analysis of the voltage dynamics acquired in (a). The electrical stimulation artifact was measured in the presence of TTX and subtracted from the action potential waveforms. n = 75 spikes for each ROI. **c**, The resulting waveforms for the 23 ROIs. **d**, Quantification of kinetics parameters from the action potential waveform. The action potential waveform and its time derivative are shown. t_FWHM_, full-width-half-maxima of the action potential. t_rise_, the 10-90% rise time. t_fall_, the 90-10% fall time. **e**, Spontaneously spiking neural dynamics acquired at 5.5 kHz. The denoised traces by applying the DeepCAD-RT are overlayed with the raw data. **f**, A representative single-trial recording of action potential waveform. The data is from ROI 9 in (e). The denoised data (magenta) is overlayed with the raw data (light pink). The residual is obtained by subtracting the denoised trace from the raw trace. **g**, Submillisecond time delays of action potentials. The arrows indicate the peaks of action potentials. The ROIs 1 and 4 indicate the soma and the connected neural process, respectively.

To further analyze the analogue waveform of the measured action potentials, we temporally aligned the action potentials using the trigger input to the electric field stimulator and averaged the 75 action potentials, referred to as stimulus-triggered averaging. This process increased the SNR by a factor of ∼8. Subsequently, to eliminate the direct electric field stimulation artifact, we treated the same neurons with a sodium channel blocker, tetrodotoxin, suppressing action potential generation, and repeated the same measurement and analysis. The obtained waveform for each ROI, corresponding only to the stimulation artifact, was subtracted from the averaged action potential waveform of the ROI (**Fig. 5b**). Individual subcellular compartments showed distinct kinetic parameters of action potentials, including rise/fall kinetics and spike width (**Fig. 5c,d**).

Lastly, we applied the DeMOSAIC acquisition to quantitatively investigate a spontaneously active functional neural circuit. In-depth understanding of neural circuit function requires a faithful quantitative analysis at the level of individual action potentials, referred to as a single-trial analysis. However, waveform analyses on individual spikes at submillisecond-scale temporal resolution (5.5 kHz) suffered from low SNR, caused primarily by shot noise. To address this issue, we employed a deep learning-based statistically non-biased denoising technique, DeepCAD-RT, to the DeMOSAIC data^30–33 44,46^ (**Fig. 5e**). To our surprise, the denoising algorithm demonstrated highly effective in suppressing the shot noise, resulting in SNR improvement of over 2-fold without introducing significant waveform distortion (**Fig. 5f**). Since single-trial data faithfully recapitulated the action potential waveforms, we were able to reliably quantify the peaks and widths of individual spikes, as well as their submillisecond-scale delays (**Fig. 5g**).

## Discussion

We have reported a novel optical segmentation-based detection scheme, DeMOSAIC, which assigns each user-defined ROI to a minimal number of pixels in the detector. By minimizing the number of used pixels in the detector, the DeMOSAIC system provides optimally compressed data acquisition, advantageous especially for high-speed functional imaging on spatially entangled structures. We demonstrate its unique detection capabilities in a synthetic graffiti sample at >100 kHz and in neuronal circuit dynamics at >5 kHz, which is challenging for the conventional detection scheme. We expect the DeMOSAIC acquisition will open new opportunities for investigating circuit-scale neural dynamics at unprecedented spatiotemporal resolutions.

In conventional ROI-based functional imaging, spatial resolution is required for defining the margins of the ROIs. Since the sensors of conventional cameras are composed of an array of square pixels, a large number of pixels is required to represent the margins and inner areas of the ROIs at high spatial precision. Conceptually, our DeMOSAIC system resolves this inefficiency in spatial representation by optically transforming the square pixel into the shape of the ROI. Employing adaptive optics, the pixel-to-ROI transformation is flexibly configured to match the spatial distribution of the ROIs. This spatial compression constitutes the essence of the DeMOSAIC acquisition for maximizing the temporal information.

The current DeMOSAIC system has room for further technical improvement. First, we used a 3-by-3 MLA for relaying the zeroth and first-order diffractions therefore the higher-order diffractions were discarded, and the collection efficiency was limited to ∼70%. Extending the MLA to 5-by-5 or larger will enable us to capture the residual higher-order signals to improve the collection efficiency (**Extended Data Fig. 4c**). Second, the field-of-view of the current DeMOSAIC system is limited by the size of SLM active window (**Extended Data Fig. 9**). Reducing the relay magnification on the SLM increases the field-of-view but also compromises the resolution for optical segmentation. Introducing the SLM with a higher pixel resolution can be a solution to this problem. The smaller pixel size also accompanies increase in the diffraction angle, thereby improving the maximum collection NA (**Extended Data Fig. 2c**). Third, the current DeMOSAIC system, designed for the near-infrared window, lacks compatibility with visible fluorophores due to the wavelength dependency of the diffraction angle (**Extended Data Fig. 1D**). The introduction of differently sized MLAs mounted in a stepper wheel could facilitate the use of diverse fluorophores across a broader spectral range. Fourth, *post hoc* computational source separation algorithms^27,34,35^, which rely on dense spatial information, may not be applicable to the data acquired by DeMOSAIC. Developing a new algorithm tailored for the spatially compressed data requires further investigation. Lastly, the ROIs are preselected for the DeMOSAIC acquisition, so that the sample needs to be stationary during the acquisition. This restriction may be ameliorated by introducing the closed-loop algorithm to update the ROI information based on intermittent high-resolution image acquisition. Alternatively, a single ROI may be divided into four subimages to function as a quadrant cell photodetector for tracking the center-of-mass while recording its intensity trace^7^.

Our DeMOSAIC system has a modular design so that it can be flexibly integrated into a camera port of various widefield microscopes (**Extended Data Fig. 10** and **Supplementary Table 2**). For example, the DeMOSAIC system may be combined with structured illumination microscopy to attain higher resolution in defining the ROIs^36,37^, or with light-sheet illumination to observe thick biological specimens such as organoids, brain tissues, and whole organisms^36–38^. Alternatively, targeted photostimulation combined with optogenetic actuators will allow precise control of neural activities while recording the circuit-scale neural dynamics^12,38–43^. Moreover, the DeMOSAIC system is also compatible with various detectors for versatile applications. Faster detectors such as a silicon photomultiplier array with a GHz digitizer can be adopted for allowing fluorescence lifetime imaging^44,45^ or fluorescence correlation spectroscopy^46,47^. We anticipate that our DeMOSAIC system will be broadly adopted to study complex dynamic phenomena.

---

**Extended Data Fig. 1:**
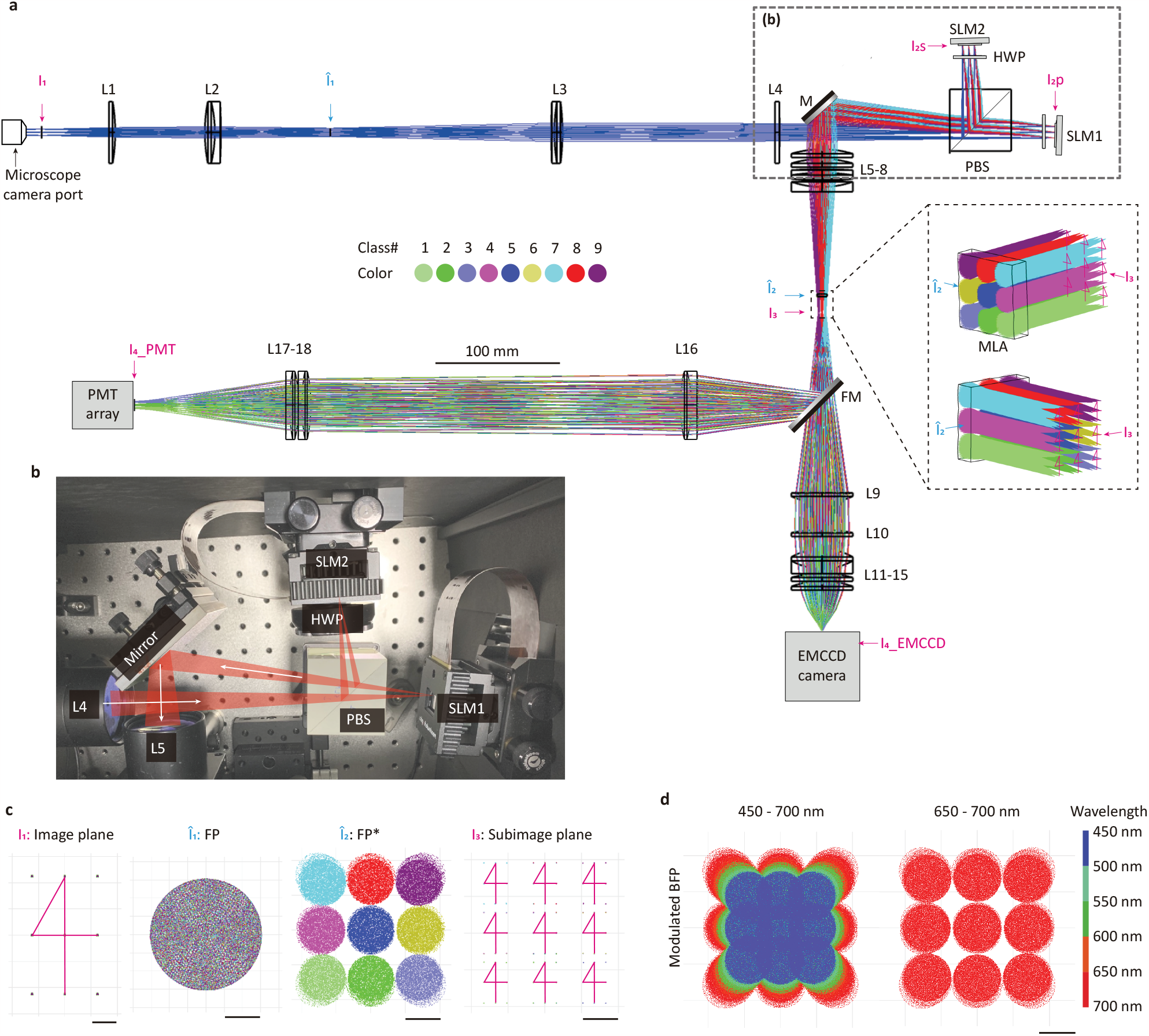
Detailed design of the DeMOSAIC system. **a**, Overall design of the DeMOSAIC system using a ray simulation in the Zemax OpticStudio software. The image plane (I_1_) at the camera port was relayed to the SLMs via L1-L4 (magnification = 1.87x). Angular diffraction towards 9 directions was simultaneously displayed. The segmented subimages (I_3_) are formed by the 3-by-3 MLA. The subsequent optical relays (L9-L15 and L16-L18) are designed to match the sensor size of EMCCD and PMT arrays, respectively. **b**, The photograph of the polarization-insensitive SLM configurations. **c**, Representative full-field spot diagrams. I_1_, 3 × 3 spots are launched at I_1_. ‘4’-shaped line was overlayed for enhancing visibility. î_1_, Fourier plane of I_1_. î_2_, Fourier plane after 8-directional angular modulations. I_3_, Image plane formed by the MLA. **d**, Wavelength dependence of the Fourier plane image after the angular modulation by SLMs (î_2_).

**Extended Data Fig. 2:**
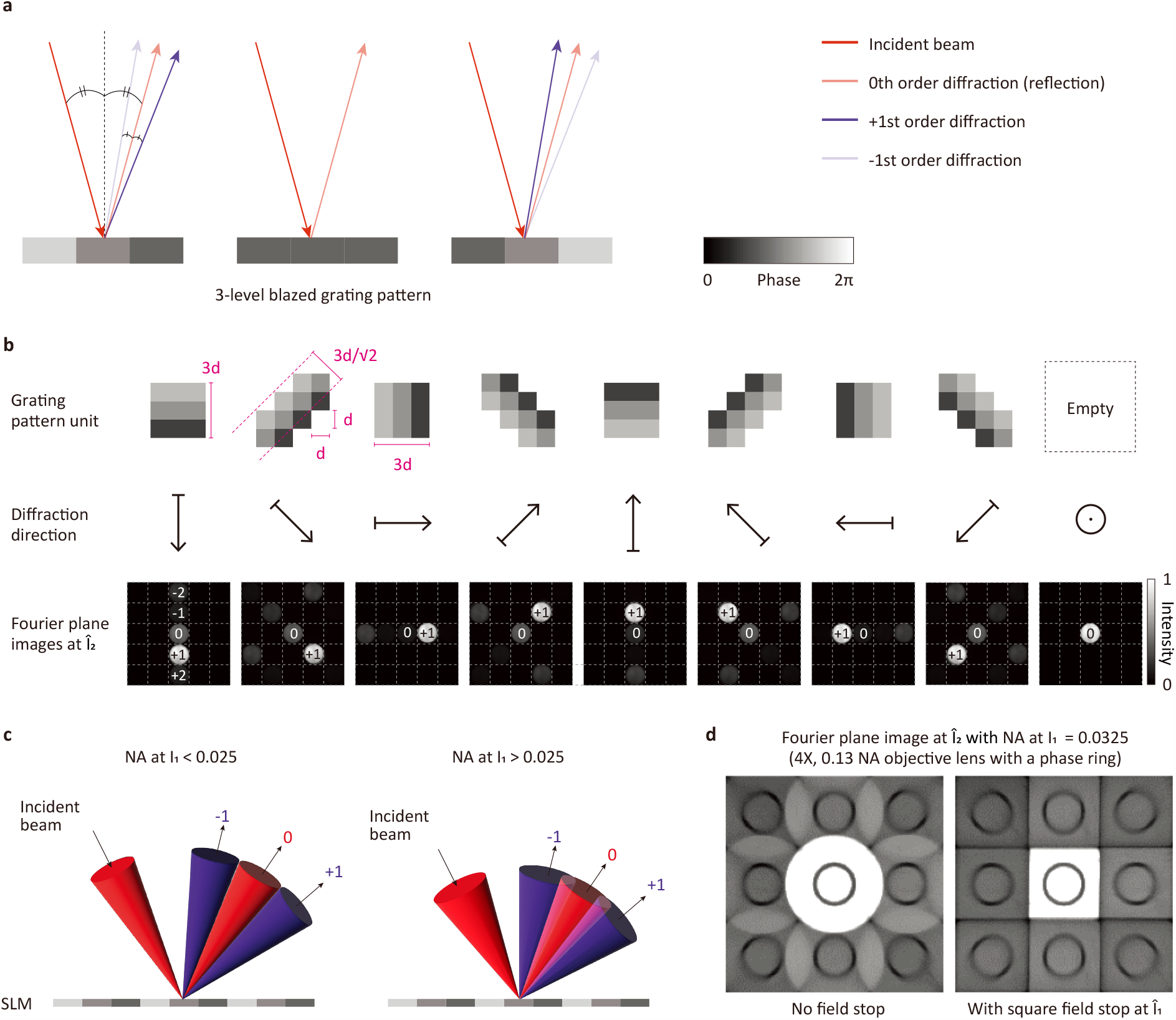
The blazed phase gratings for the spatial light modulator. **a**, Schematic illustration of the diffraction by the 3-level blazed phase grating pattern displayed on SLM. The higher-order diffractions are omitted. **b**, The grating pattern units for 9 directions and their corresponding Fourier plane images at the î_2_ plane. The Fourier plane images are taken by placing a CCD camera at the plane where the MLA is placed. **c**, Incident NA dependent diffraction crosstalk. Our DeMOSAIC design supports the incident NA of up to 0.025. If the incident NA is larger than 0.025, there is an overlap among nearby diffraction orders. **d**, Field stop. To avoid the crosstalk among nearby diffraction orders, a square-shaped field stop is placed

**Extended Data Fig. 3:**
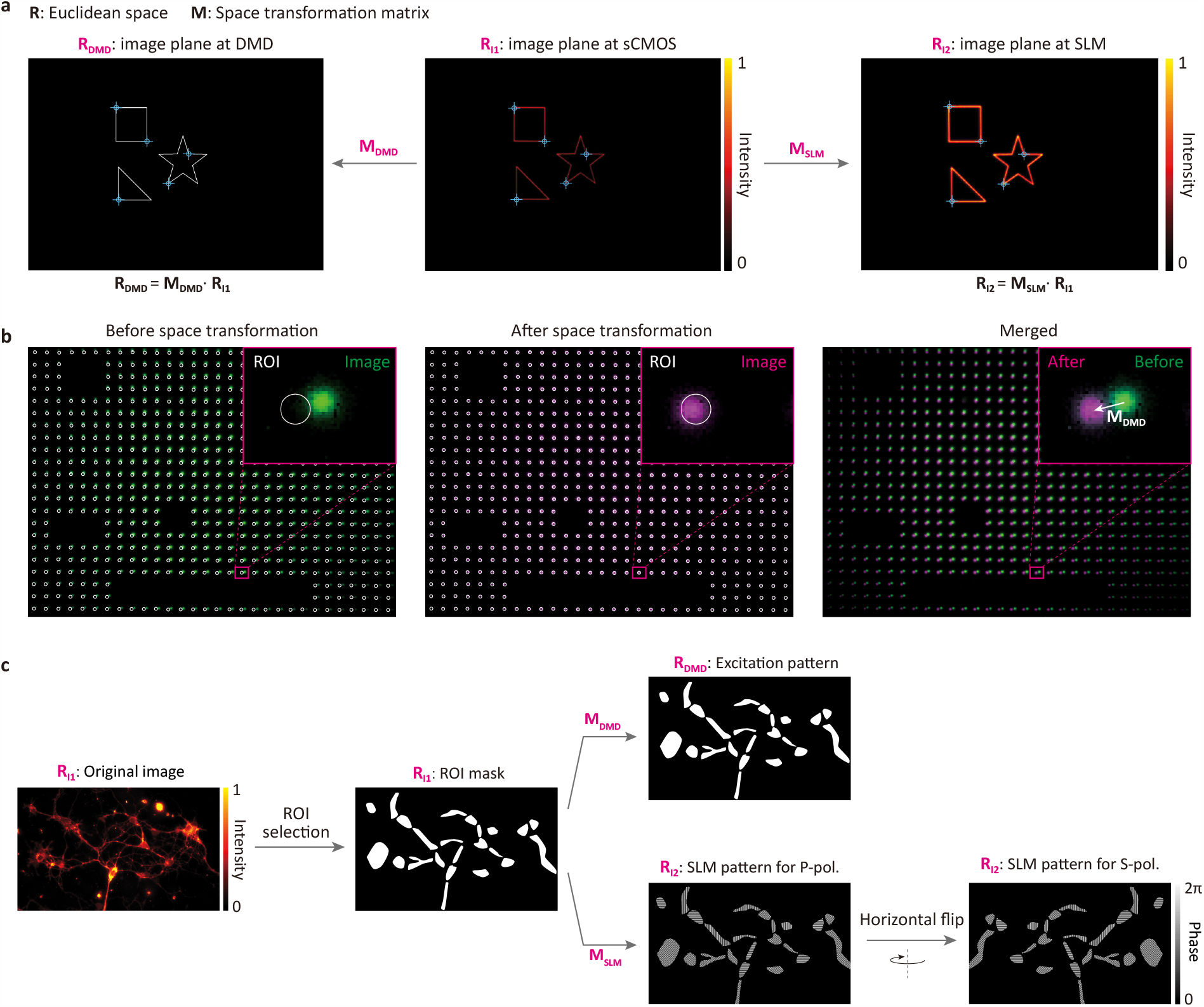
Coregistration of conjugated image planes. **a**, Obtaining space transform matrices using a point-based multimodal image coregistration algorithm. We generated arbitrary simple patterns for DMD and projected light pattern on the fluorescence dye coated mirror surface at the objective lens focal plane. The projected image was relayed to both sCMOS camera plane (I_1_) and SLM plane (I_2_). By comparison of relative coordinates of manually selected points, we could obtain transform matrices between spaces. M_DMD_ and M_SLM_ refers transform matrices from the sCMOS camera space to DMD and SLM, respectively. **b**, Demonstration of the coregistration. The mismatch between the selected ROI in the original image plane and the image plane at the SLM is corrected by applying the space transformation matrices acquired in (a). **c**, Generation of ROI based patterns. A binary ROI image obtained from image space (R_I1_) was transformed to M_DMD_ and M_SLM_ matrices into DMD space (R_DMD_) and SLM space (R_I2_), respectively. Patterns for SLMs and DMD2 (for optogenetic stimulation pattern) are prepared by horizontal flipping of paired pattern images Demonstration of optical segmentation.

**Extended Data Fig. 4:**
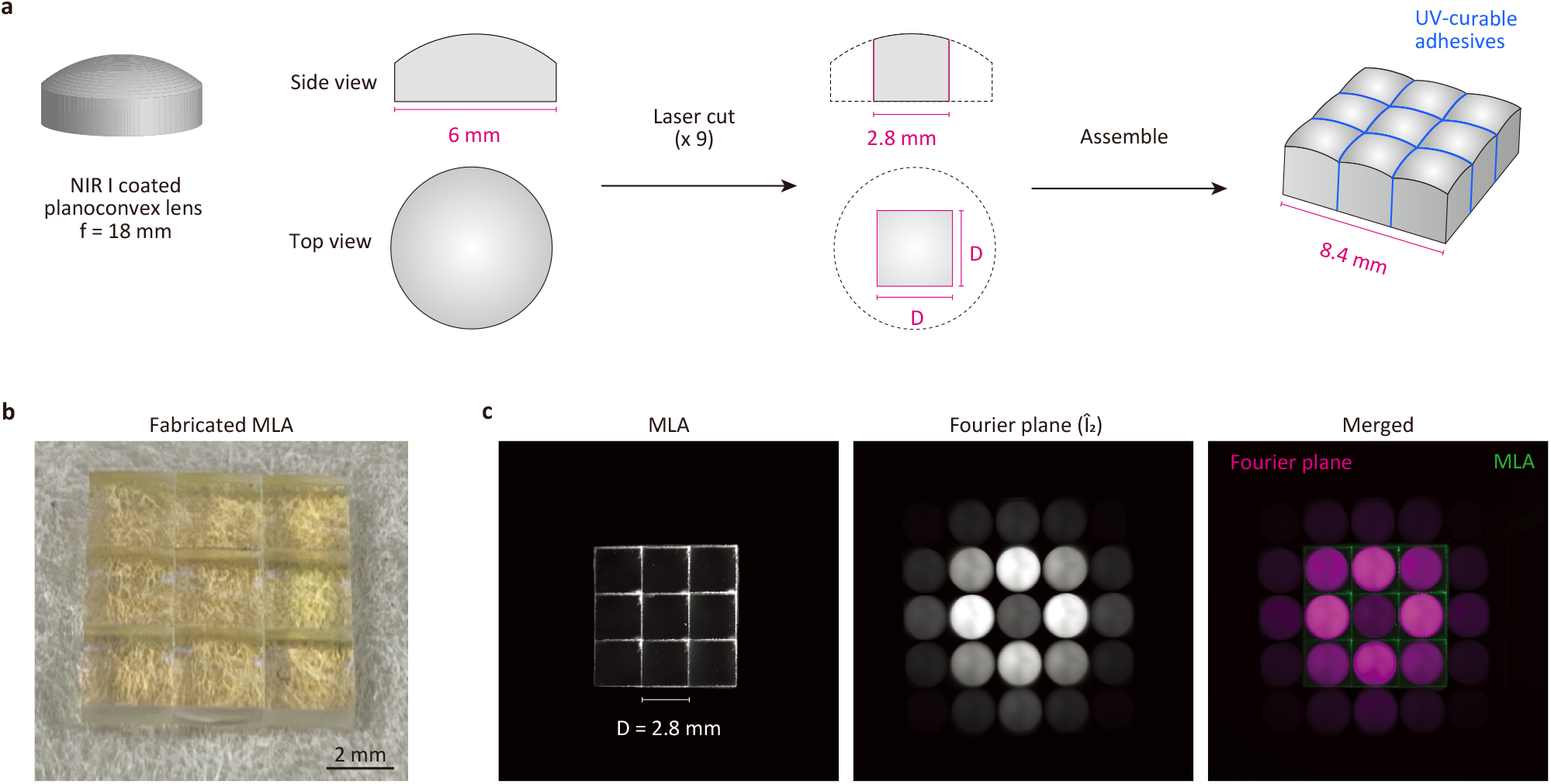
Fabrication of the microlens array (MLA) **a**, The fabrication procedure. Plano-convex lenses with NIR-I antireflection coating were cut into a square shape and assembled to a 3 by 3 grid with an optical adhesive. **b**, The photograph of the fabricated MLA. **c**, Alignment of MLA and the Fourier plane image at î_2_. The Fourier plane image represents the maximum intensity projection of all the 8-directions. Note that the conjugated back aperture plane for each subimage is aligned with each lenslet in the MLA. The second order diffractions surrounding the MLA were blocked.

**Extended Data Fig. 5:**
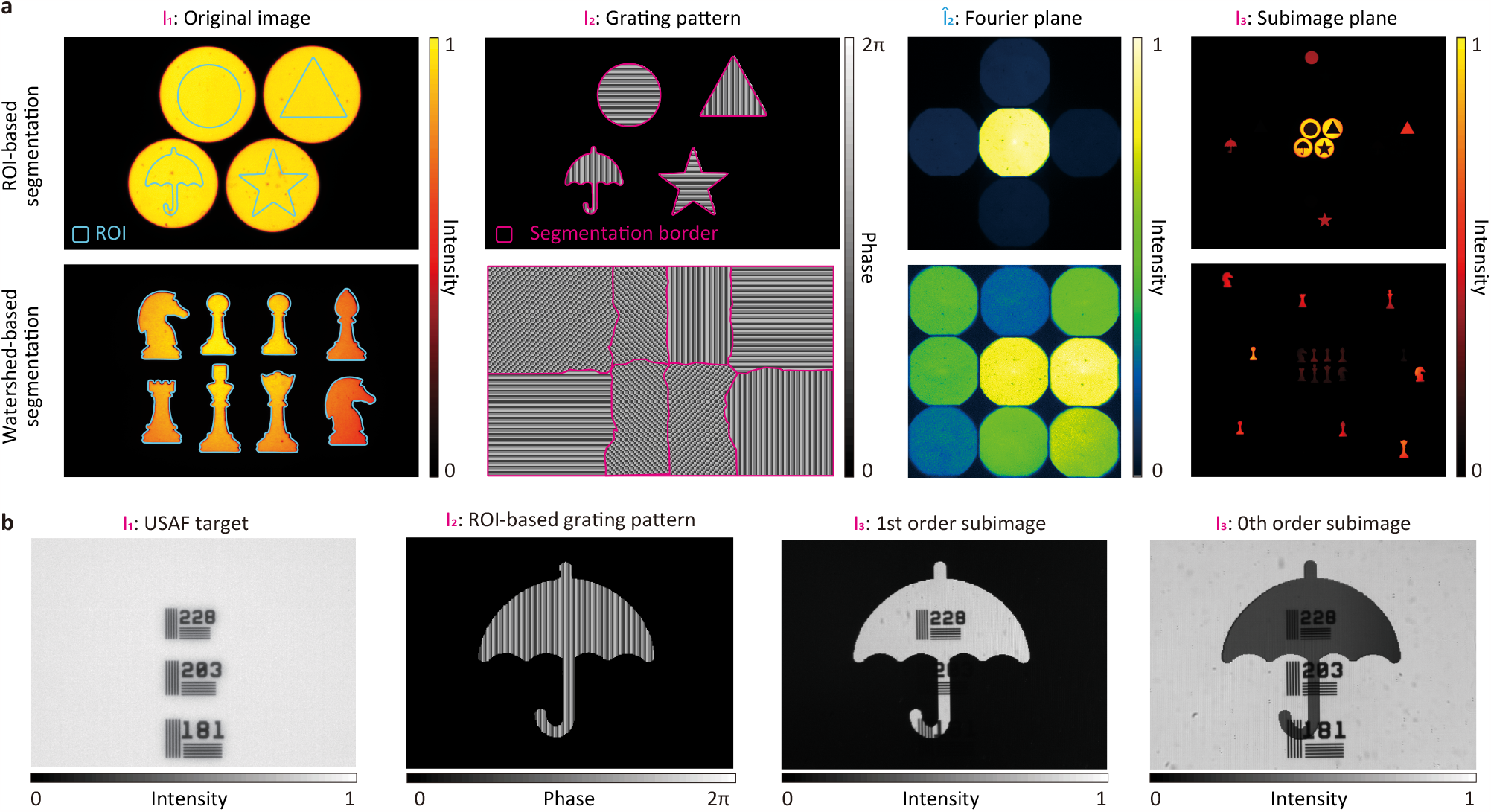
Demonstration of optical segmentation by the DeMOSAIC system. **a**, The two algorithms of generating the grating patterns. The upper row shows the ROI-based grating pattern, which can be generally used for segmenting the ROIs. In this case, the segmentation border matches with the ROIs. If the ROIs exclusively have the signal, as in the lower row, the watershed algorithm can be applied. The watershed algorithm was applied combined with patterned excitation targeted on the ROIs. **b**, Demonstration of optical segmentation of a reflectance image. The reflectance image of the USAF target was segmented in an umbrella shape. The +1st order and 0th order subimages show mutually exclusive patterns. Note that the internal pattern in the +1st order subimage is preserved after optical segmentation.

**Extended Data Fig. 6:**
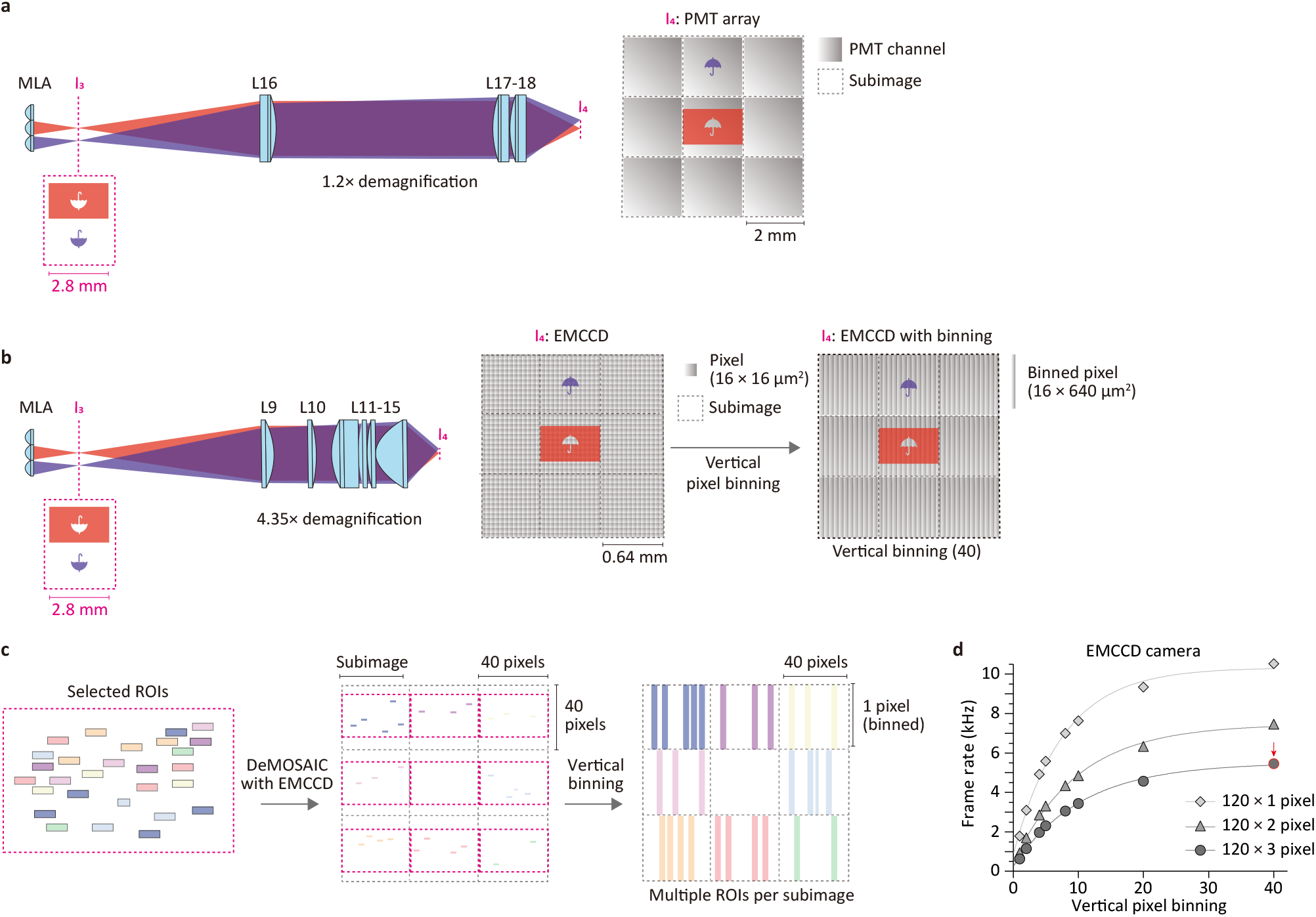
Detector configurations. **a**, Optical relay for the PMT array. To match the sizes of a subimage and a PMT channel, the optical relay with lenses, L16-L18, demagnified the image by a factor of 1.2. In this configuration, each ROI is assigned to a PMT channel. **b**, Optical relay for the EMCCD camera detector. The optical relay (L9-L15) is designed to have a demagnification factor of 4.35, projecting the 3×3 subimages onto 120 × 120 pixels in the EMCCD. The vertical pixel binning of 40 pixels is applied, reducing the pixel dimension to 120 × 3 pixels. **c**, Schematic representation of assigning multiple ROIs on individual subimages. **d**, The measured frame rate of the EMCCD (iXon Ultra 897, Oxford Instruments) with different pixel dimensions. The arrow indicates the parameter used in this study, corresponding to the frame rate of 5.5 kHz at 120 × 3 pixels.

**Extended Data Fig. 7:**
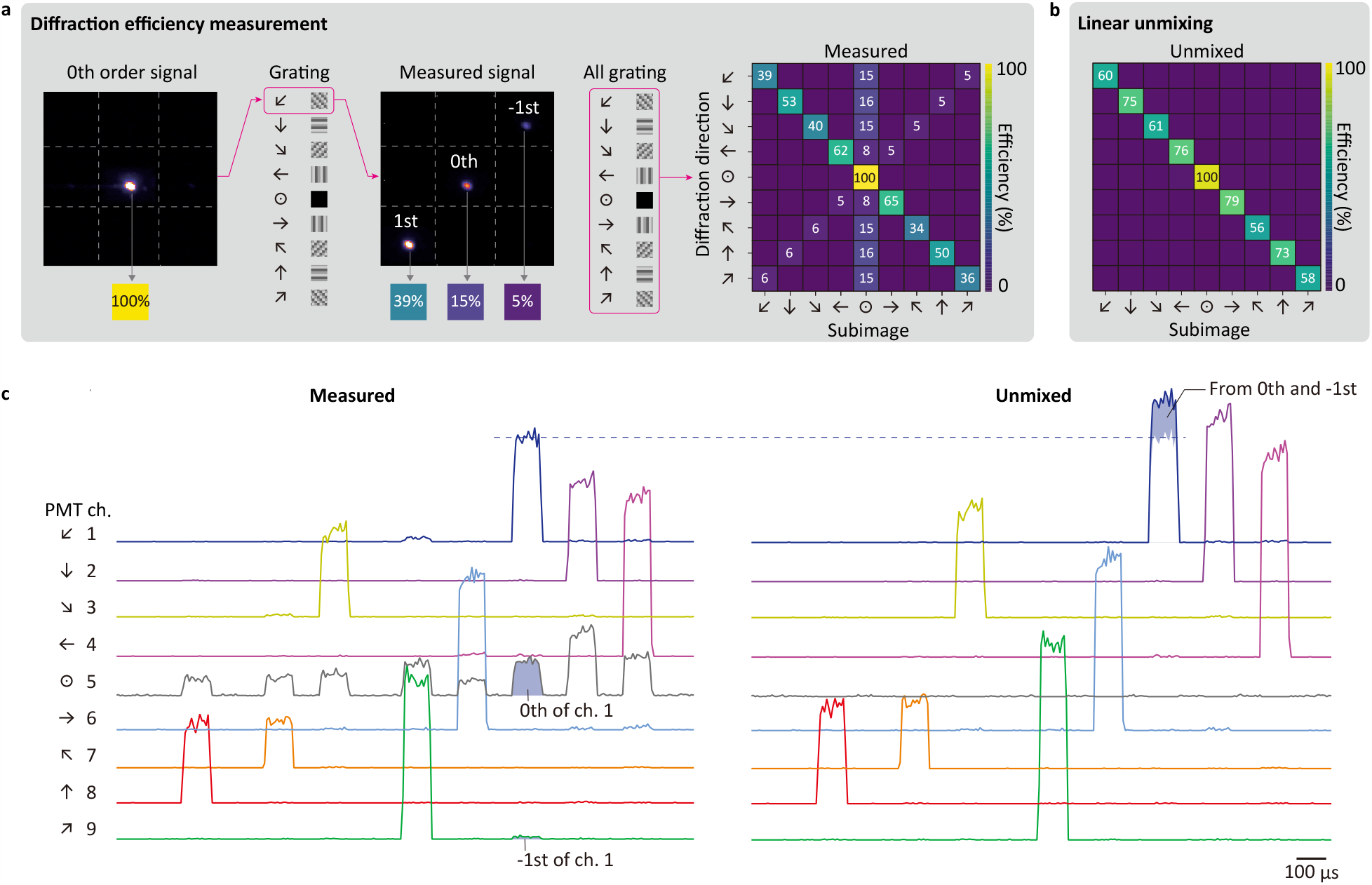
Linear unmixing of diffraction crosstalk. **a**, The measured first-order diffraction efficiency. The efficiencies were normalized to the zeroth-order signal measured without the grating pattern. **b**, Linear unmixing. The 0th and −1st order signals are combined with the corresponding 1st order signals. The mean calibrated efficiency was ∼67%. The losses are mostly attributed the ±2nd orders. **c**, Demonstration of linear unmixing on the experimental data in Figure 3. The channel 5 corresponding to the zeroth order was removed after applying the linear unmixing. The shaded regions in the ‘measured’ traces (left) represents the 0th and −1st order signals for the channel 1. These signals are moved to the channel 1 in the ‘unmixed’ traces (right).

**Extended Data Fig. 8:**
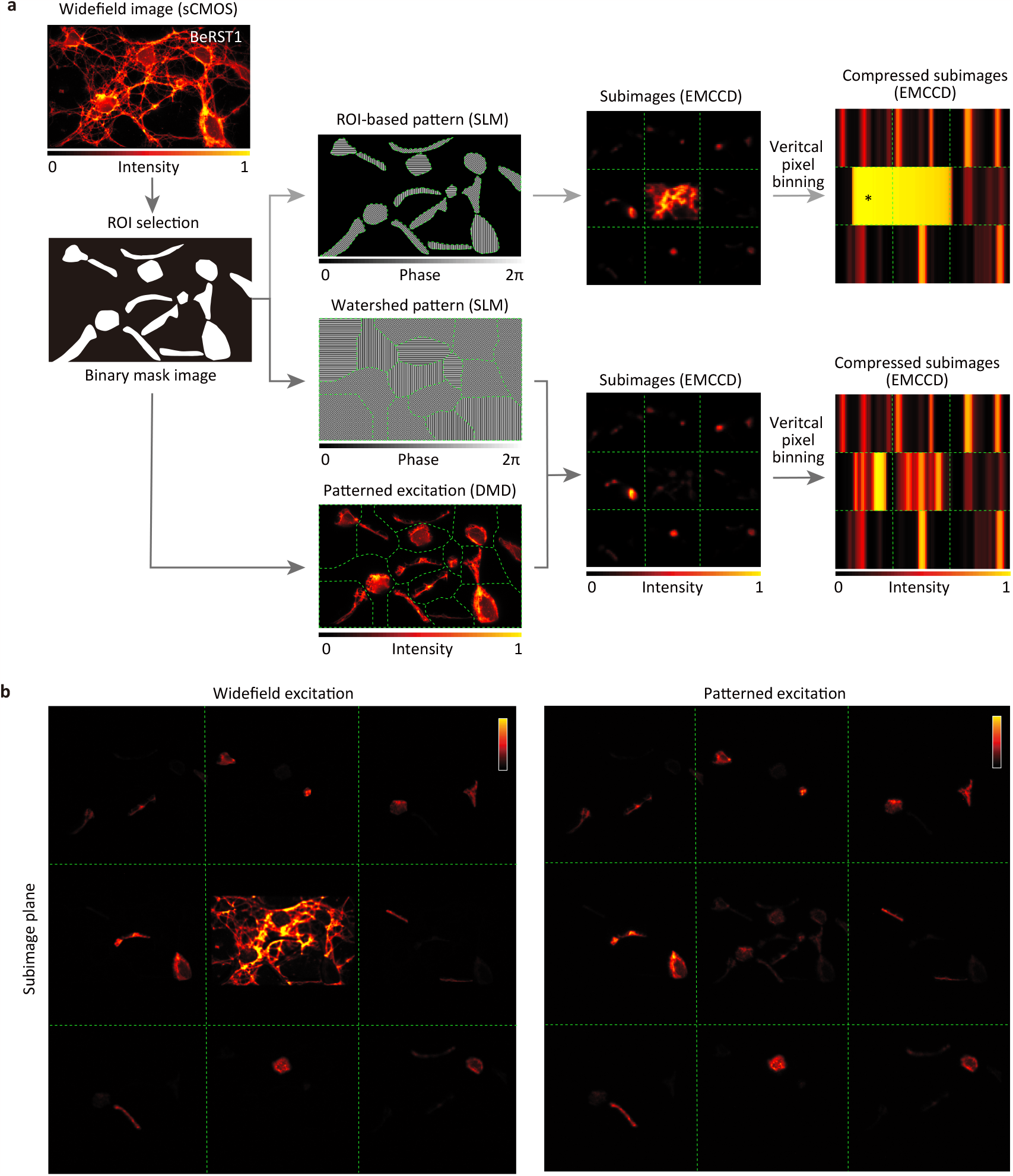
Patterned excitation with DeMOSAIC. **a**, The DeMOSAIC acquisition pipeline with patterned excitation. The binary image of the selected ROIs is used to create the patterned input to the DMD and to generate the phase grating using the watershed algorithm. Note that the zeroth-order signal is dramatically reduced when patterned excitation is applied. The asterisk indicates pixel crosstalk from the saturated zeroth-order signal. **b**, The magnified view of the subimage planes with widefield excitation (left) and patterned excitation (right).

**Extended Data Fig. 9:**
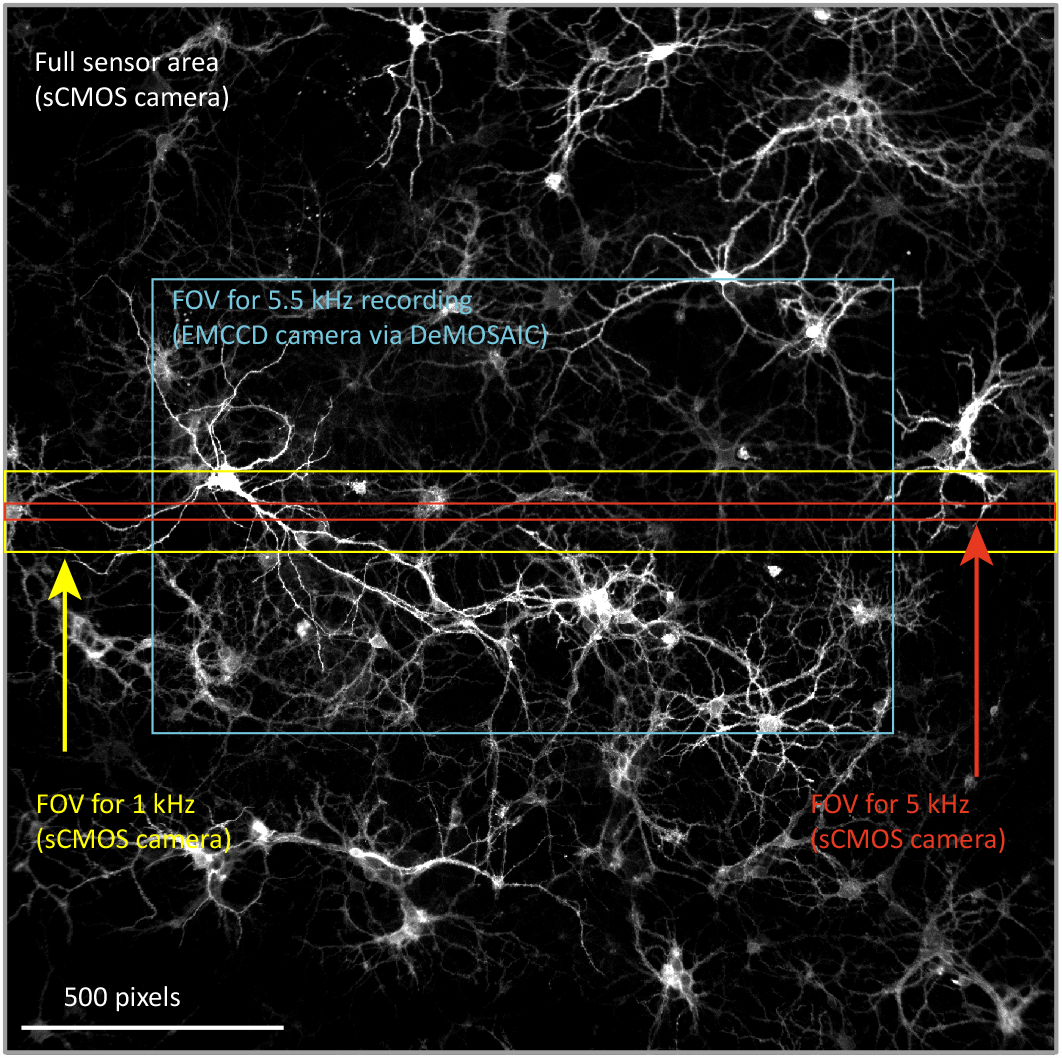
Comparison of the field-of-views (FOV). The effective FOV for conventional widefield imaging system equipped with an sCMOS camera and the DeMOSAIC system equipped with an EMCCD camera was shown. The effective FOV of our DeMOSAIC acquisition was ∼30% of the full FOV of the conventional C-mounted sCMOS camera. The larger FOV can be obtained by introducing a larger SLM. Alternatively, the magnification factor of the optical relay to the SLM may be decreased by compromising the resolution of optical segmentation. In contrast, a conventional sCMOS camera provides a frame rate of 1 kHz only by reducing the FOV down to 7.7% by subarray readout. Note that FOV can be varied by camera model and the background neuron image was taken by a 10x objective lens only for the visualization purpose.

**Extended Data Figure 10:**
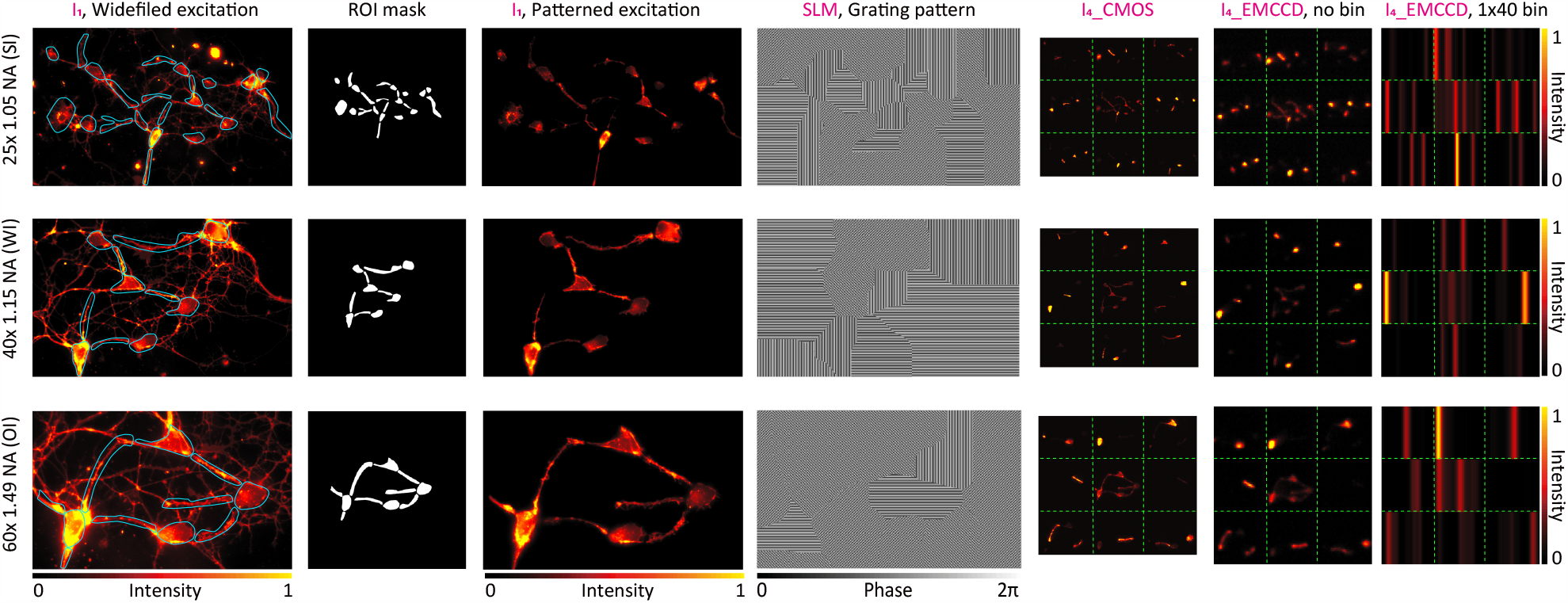
Demonstration of the DeMOSAIC system with different objective lenses. Our DeMOSAIC system is compatible with equipping various types of objective lenses. The figure shows the representative 3 types of objective lenses with different magnifications, NA, and immersion media.

## Methods

### System design

The DeMOSAIC system was designed based on the ray tracing simulation in the Zemax OpticStudio (**Extended Data Fig. S1**). The source rays (650–700 nm) were launched from an original image plane at the sample (I_1_) with an imaging NA of 0.025 (i.e., objective NA divided by the magnification). The image plane (I_1_) is relayed by a series of lenses (L1-L4) to the conjugated image planes (I_2_), where a pair of reflective SLM is placed. Each SLM was simulated as a diffraction grating with pixelated phase steps (pixel size = 9.2 μm), and is tilted by 6º for securing the minimal angle of reflection of 12º toward the detector path. The SLM plane is then relayed to the subimage plane (I_3_) formed by a 3 × 3 MLA, which was custom-built to match the size of the conjugated BFP image (î_2_). The subimage plane was magnified or demagnified to match the detector sensor sizes (magnification factor: 0.23x for the EMCCD and 0.82x for the PMT array). In addition to the optical parts, we considered dimensions of optomechanical components to confirm feasibility of implementing the designed system.

### Implementation of the DeMOSAIC module

According to the optimized design from the ray tracing simulation, we built the DeMOSAIC system on the inverted epifluorescence microscope equipped with a motorized filter wheel and two camera ports (Eclipse Ti2, Nikon; **Extended Data Fig. 1**). NIS-Elements software (Nikon) was used for controlling the filter wheel and cameras. On the left-side camera port, we mounted an sCMOS camera (ORCA-Flash 4, Hamamatsu) providing a snapshot image of widefield reflectance or fluorescence for user-defined selection of the ROIs. A motorized flip mirror was used to redirect the beam path to the rightside camera port where the DeMOSAIC module is implemented. The image plane at the sample (I_0_) was relayed to the conjugated image plane at an SLM (I_2_^s^ and I_2_^p^) via L1-L4 with 1.87x magnification. A pair of SLMs receives input of a ROI-based blazed grating pattern and introduces phase modulation for image segmentation (**Figure 2a**). To block the relay of unwanted reflected beam from the outside of active pixel window of SLM, a rectangular-shaped aperture stop was placed at the intermediate image plane (I_1_). Due to the polarization sensitivity of SLM, unpolarized fluorescence emission was split into linear polarized beams via a polarizing beam splitter, which were modulated by two SLMs, one for s-polarization (I_2_^s^) and the other for p-polarization (I_2_^p^). Since we set the orientation of the SLM to p-polarization, a half-waveplate (WPH10E-670, Thorlabs), introducing 90º polarization rotation, was placed in front of the SLM in the s-polarized beam path (I_2_^s^). By adjusting the position and tilt of the SLM mounts, the image planes for the two SLMs were coaligned in a sub-pixel precision. The reflected beams from the SLMs were redirected by a mirror to pass through a set of lenses (L5-L8) and reaches the custom-built 3 × 3 MLA positioned at the second-conjugated back-focal-plane (î_2_) to form a subimage image plane (I_3_). A rectangular-shaped aperture stop was placed at the first conjugated pupil plane (î_1_) to block inter-class spill-over at the MLA (**Extended Data Fig. 2**).

Segmented subimages (I_3_) were relayed to one of two types of detectors (I_4_), an EMCCD camera (iXon Ultra 897, Oxford Instruments) and a 64-channel PMT array (H7546B-20, Hamamatsu). The optical path to each detector was switched by beam-turning mirror cubes (DFM2RM, Thorlabs). The relay to the EMCCD was with 0.23x magnification (i.e., 4.35x demagnification) to minimize the number of pixels to 120 × 120 pixels for high-speed readout with ‘isolated crop mode’. For the PMT array, the relay system is designed as 0.82x magnification (i.e., 1.2x demagnification) to match the center-to-center pitch of subimages (2.8 mm) to that of the PMT array (2.3 mm). The relay system to each detector is designed to minimize system aberrations (e.g., spherical aberrations from the MLA), resulting in near diffraction-limited optical performance.

### Co-registration of conjugated image planes

Spatial coordinates of conjugated sCMOS camera sensor space (R^I1^), DMD space (R^DMD^), and SLM space (R^I2^) are coregistered by applying linear transformation matrices. First, a mirror surface was placed at the sample plane of the objective lens to obtain a transformation matrix (M_DMD_) between the sCMOS camera and the DMD. An arbitrarily shaped pattern generated by the DMD (e.g., star) was projected on the mirror surface and a reflectance image was acquired by an sCMOS camera. The transformation matrix was obtained by applying a point-based coregistration algorithm between the DMD input and the image acquired from the sCMOS using MATLAB (MathWorks). Using an auxiliary CMOS camera (temporally used only for coregistration) conjugated to the SLM window (I_2_), the same procedure was performed to obtain a transformation matrix (M_SLM_) between the sCMOS camera and the SLM. By applying the acquired transformation matrices, an excitation pattern, and a pair of segmentation grating patterns from the binary image of selected ROIs were generated. In our implementation, the transformation matrices were nearly invariant over 6 months. The MATLAB codes are available on the GitHub repository.

### Generation of ROI-based phase grating pattern

Grey values for 3-level pixelated grating were determined from phase retardation and grey value calibration data provided for each SLM head. The steering angle for the m-th order (*θ*_*m*_) by pixelated grating is determined using the following relation:

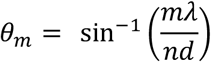

where *λ* is wavelength of incident light, *d* is pixel pitch and *n* is a step of blazed grating. In our design, 3-level and 8-bit grayscale values were used to construct the 8-directional blazed grating pattern units. The phase grating pattern image for the SLM^p^ was generated by filling a desired grating pattern unit in each ROI of an ROI-based binary image via MATLAB codes. The corresponding input for the SLM^S^ was created by horizontally flipping the phase grating pattern image for SLM^P^. The MATLAB codes are available on the GitHub repository.

### A sequential light pattern generation of dynamic graffiti

The 8 binary digital images for each letter of the graffiti on ‘DEMOSAiC’ were generated for demonstration purpose and loaded in the internal memory of DMD coupled with red LED (Solis 660C, Thorlabs). A temporal sequence of each image is assigned to form a message ‘i AM CODES’ with a frame on time 105 μs and off time 105 or 210 μs, respectively (**Fig. 3d**). The light image pattern was reflected by a beam splitter (BSW29R, Thorlabs) and projected to the mirror at the sample plane, which was conjugated to the DMD. The returning beam was relayed to DeMOSAIC acquisition beam path equipped with a 64-ch PMT array.

### DeMOSAIC acquisition at 125 kHz with PMT array

The PMT array was constructed with a 64-channel multianode photomultiplier tube assembly (H7546B-20, Hamamatsu) connected to the interface board (SIB164B, Vertilon) and a 64-channel pulse counting system (MCPC682, Vertilon). Acquisition parameters of the PMT array was controlled by the vendor-supplied software (Control and Acqusition Interface, Vertilon). High voltage and pulse threshold were set to −920 V and 7 mV, respectively. The sampling rate was set to 125 kHz, which was limited by the minimal charge integration interval of the 64-channel pulse counting system (8 μs). Among 64 channels, 9 channels (3 x 3 array) were used for recording the magnified subimage plane.

### Primary neuron culture

All animal procedures were approved by the Seoul National University Institutional Animal Care and Use Committee and complied with all relevant ethical regulations for animal research (IACUC #SNU-220616-1-1). Dissociated neurons were harvested from dissected hippocampi of wild-type Sprague Dawley rat pups at embryonic day 17-18. The cells were plated at a density of 100 cells per mm^2^ on an 18 mm coverslip coated with poly-D-lysine and laminin (Neuvitro, Vancouver, WA) in a culture medium containing neurobasal growth media (NbActiv4, BrainBits), 1% penicillin-streptomycin (P4333, Merck), 1% N-2 supplement (17502048, ThermoFisher Scientific), 10 ng/ml of brain-derived neurotrophic factor (BDNF; 248-BDB-010, R&D Systems), and 10 ng/ml of glia cell line-derived neurotrophic factor (GDNF; 212-GD-010, R&D Systems). The cells were placed in an incubator maintained at 37*°*C and 5% CO_2_. Every 3-4 days, half of culture media was replaced to the fresh media. Cultured neurons were used for optical imaging between days in vitro 9 and 14.

For calcium imaging, the culture medium was exchanged to 1 μM Cal630-AM (20721, AAT Bioquest) dissolved in 10% Pluronic F-127 with imaging solution containing 140 mM NaCl, 10 mM HEPES, 30 mM glucose, 3 mM KCl, 1 mM MgCl_2_ and 2 mM CaCl_2_ with the pH adjusted to 7.3 with NaOH. The neurons were incubated in an incubator for 2 hours and then washed twice with fresh imaging solution. For voltage imaging, the neurons were incubated with 1 μM BeRST1 dissolved in imaging solution for 20 min, and washed twice with fresh imaging solution. The stained neurons were mounted in a homeothermic imaging chamber (CMB-18-EC-PB, Live Cell Instrument) for data acquisition.

### DeMOSAIC acquisition of neuronal dynamics at 5.5 kHz with EMCCD

For imaging neuronal dynamics with DeMOSAIC system, a widefield fluorescence image of BeRST1 was captured by an sCMOS camera. From the snapshot image, multiple ROIs were defined by using the manual ROI selection function in NIS-Elements software, resulting in the binary images of ROIs. The ROI-based grating patterns, as well as a corresponding binary image for patterned excitation, were generated by MATLAB codes and displayed to the SLMs and the DMD. For excitation of BeRST1 dyes, we coupled 620 nm wavelength light source (Solis-620D, Thorlabs) to DMD and the excitation power did not exceed 10 mW/mm^2^ in this study^48^.

For achieving the maximal acquisition speed with the ‘isolated crop mode’, the conjugated image plane is aligned to be detected at the lower-right corner of 120 x 120 pixels in EMCCD camera (iXon Ultra 897, Oxford Instruments). The camera was controlled using the Andor SOLIS software. The camera was cooled to −80 ºC prior to imaging using a recirculating water chiller (UC160-190, Solid State Cooling Systems). Acquisition parameters were set to the following: vertical pixel shift of 0.3 μs, vertical clock voltage amplitude of +2, readout rate of 17 MHz, bit-depth of 16-bit, electron multiplier (EM) gain of 150, and pre-amplifier gain of 1x. In addition, asymmetric vertical binning of 40 pixels was applied to achieve a frame rate of 5.5 kHz at 120 × 3 pixels. Typically, we acquired 180,000 frames for 32 s for 5.5 kHz which is saved in a TIFF file of 174 MB. For the calcium imaging study, we set the frame rate to 200 Hz considering the slow kinetics of the intracellular calcium signals.

Temporal sequence of electrical field stimulation and camera acquisition trigger was controlled by a multichannel pulse stimulator (Master-9, AMPI).

## Data analysis

Voltage imaging data acquired at 5.5 kHz is saved as a 16-bit TIFF file with 120 × 3 × 180,000 pixels (120 × 3 pixels, 180,000 frames, 174 MB). The raw data was first processed for unmixing the crosstalk between +1 and −1 order diffractions (**Supplementary Section 2**). The unmixed data was then denoised for spontaneous activity data by DeepCAD-RT. The parameter set for the training for DeepCAD-RT^30^ were ‘patch_xy = 10’, ‘patch_t = 1000’, and ‘epoch = 20’. The training typically took∼6 hours in a PC equipped with two GTX3080 GPUs. Time-series intensity trace for each ROI was extracted and is represented as dF/F_0_ by normalizing with the baseline fluorescence. The baseline fluorescence was measured from the 0.49 quantile values for each 1000-frame sliding window (−499 to +500 frames) using a customized python script. Calcium imaging, electrical stimulation and the graffiti data were processed by the same pipeline, excluding the denoising setup.

For stimulus-triggered averaging of electrical field stimulation evoked neuronal activity, stimulation trigger timestamp was used to align each action potential waveform, with the 400-frame time window extracted from the dF/F_0_ data, placing the stimulation timing at the 50th frame.

## Data availability

Source data are provided with this paper. All other data are available from the corresponding authors upon reasonable request.

## Code availability

All the source codes written in Python and MATLAB are available on a GitHub repository: https://github.com/Neurophotonic/DeMOSAIC

## Acknowledgments

This work was supported by the Basic Science Research Program through the National Research Foundation of Korea (NRF), funded by the Ministry of Education, Science, and Technology (2020R1A5A1018081, 2020R1A5A1018081, 2019M3E5D2A01058329, 2019M3C7A103207621) and by the Human Frontier Science Program (RGY0068/2020). We thank Seok-Hyun Yoon for supporting initiation of this work.

## Author Contributions

S.K. initiated this study. M.C. supervised the research. S.K. implemented the DeMOSAIC system and conducted experiments. J.W. and Q.D. contributed to designing the optical system. S.K., G.K., and I.K. analyzed the data. Y.L., J.W., and Q.D. contributed to applying the denoising algorithm. H.T., L-Z.F., and A-E.C. contributed to the initial conceptualization, preparation of neurons, and acquisition of preliminary data. S.K. and M.C. co-wrote the paper. All authors reviewed and edited the paper.

## Competing Interests

M.C. and S.K. are inventors of the patent-pending technology regarding the DeMOSAIC system (Korean patent application, 10-2022-0116427). No other authors declare competing interests.

**Supplementary Table 1.**
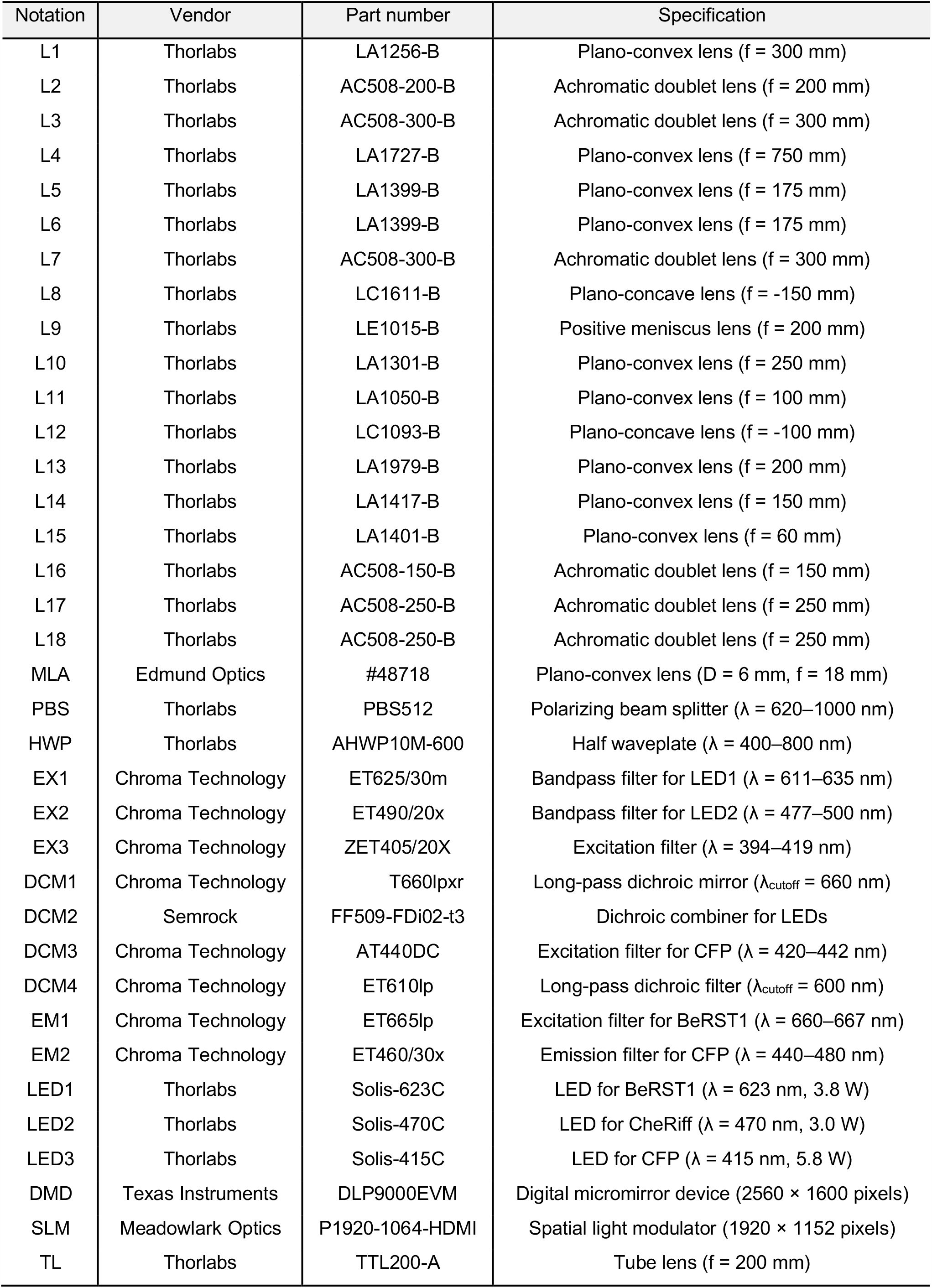
The list of optical parts used for building the DeMOSAIC system.

| Notation | Vendor | Part number | Specification |
| --- | --- | --- | --- |
| L1 | Thorlabs | LA1256-B | Plano-convex lens ($f = 300$ mm) |
| L2 | Thorlabs | AC508-200-B | Achromatic doublet lens ($f = 200$ mm) |
| L3 | Thorlabs | AC508-300-B | Achromatic doublet lens ($f = 300$ mm) |
| L4 | Thorlabs | LA1727-B | Plano-convex lens ($f = 750$ mm) |
| L5 | Thorlabs | LA1399-B | Plano-convex lens ($f = 175$ mm) |
| L6 | Thorlabs | LA1399-B | Plano-convex lens ($f = 175$ mm) |
| L7 | Thorlabs | AC508-300-B | Achromatic doublet lens ($f = 300$ mm) |
| L8 | Thorlabs | LC1611-B | Plano-concave lens ($f = -150$ mm) |
| L9 | Thorlabs | LE1015-B | Positive meniscus lens ($f = 200$ mm) |
| L10 | Thorlabs | LA1301-B | Plano-convex lens ($f = 250$ mm) |
| L11 | Thorlabs | LA1050-B | Plano-convex lens ($f = 100$ mm) |
| L12 | Thorlabs | LC1093-B | Plano-concave lens ($f = -100$ mm) |
| L13 | Thorlabs | LA1979-B | Plano-convex lens ($f = 200$ mm) |
| L14 | Thorlabs | LA1417-B | Plano-convex lens ($f = 150$ mm) |
| L15 | Thorlabs | LA1401-B | Plano-convex lens ($f = 60$ mm) |
| L16 | Thorlabs | AC508-150-B | Achromatic doublet lens ($f = 150$ mm) |
| L17 | Thorlabs | AC508-250-B | Achromatic doublet lens ($f = 250$ mm) |
| L18 | Thorlabs | AC508-250-B | Achromatic doublet lens ($f = 250$ mm) |
| MLA | Edmund Optics | #48718 | Plano-convex lens ($D = 6$ mm, $f = 18$ mm) |
| PBS | Thorlabs | PBS512 | Polarizing beam splitter ($\lambda = 620\text{--}1000$ nm) |
| HWP | Thorlabs | AHWP10M-600 | Half waveplate ($\lambda = 400\text{--}800$ nm) |
| EX1 | Chroma Technology | ET625/30m | Bandpass filter for LED1 ($\lambda = 611\text{--}635$ nm) |
| EX2 | Chroma Technology | ET490/20x | Bandpass filter for LED2 ($\lambda = 477\text{--}500$ nm) |
| EX3 | Chroma Technology | ZET405/20X | Excitation filter ($\lambda = 394\text{--}419$ nm) |
| DCM1 | Chroma Technology | T660lpxr | Long-pass dichroic mirror ($\lambda_{\text{cutoff}} = 660$ nm) |
| DCM2 | Semrock | FF509-FDi02-t3 | Dichroic combiner for LEDs |
| DCM3 | Chroma Technology | AT440DC | Excitation filter for CFP ($\lambda = 420\text{--}442$ nm) |
| DCM4 | Chroma Technology | ET610lp | Long-pass dichroic filter ($\lambda_{\text{cutoff}} = 600$ nm) |
| EM1 | Chroma Technology | ET665lp | Excitation filter for BeRST1 ($\lambda = 660\text{--}667$ nm) |
| EM2 | Chroma Technology | ET460/30x | Emission filter for CFP ($\lambda = 440\text{--}480$ nm) |
| LED1 | Thorlabs | Solis-623C | LED for BeRST1 ($\lambda = 623$ nm, 3.8 W) |
| LED2 | Thorlabs | Solis-470C | LED for CheRiff ($\lambda = 470$ nm, 3.0 W) |
| LED3 | Thorlabs | Solis-415C | LED for CFP ($\lambda = 415$ nm, 5.8 W) |
| DMD | Texas Instruments | DLP9000EVM | Digital micromirror device ($2560 \times 1600$ pixels) |
| SLM | Meadowlark Optics | P1920-1064-HDMI | Spatial light modulator ($1920 \times 1152$ pixels) |
| TL | Thorlabs | TTL200-A | Tube lens ($f = 200$ mm) |

**Supplementary Table 2.**
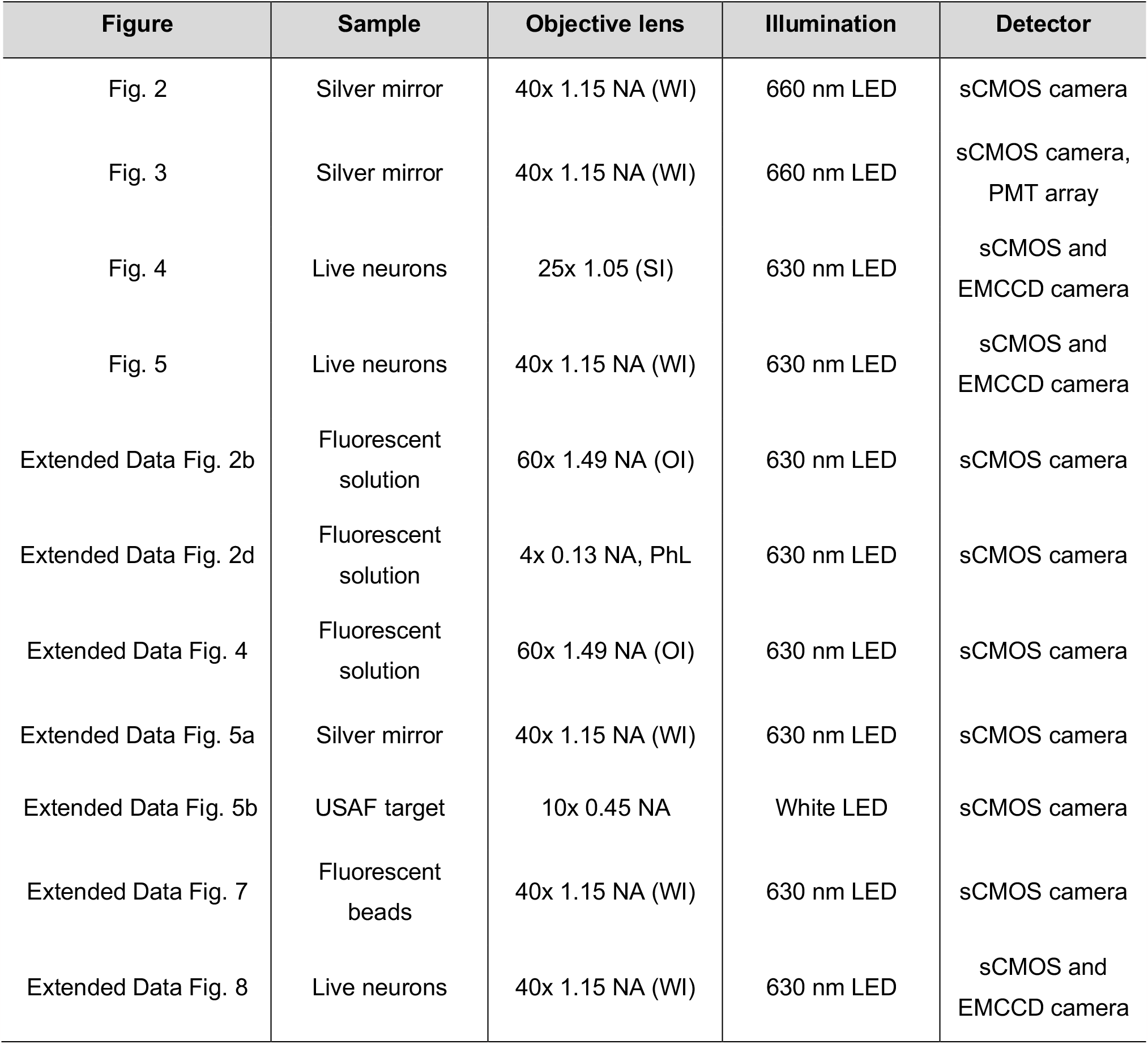
Imaging parameters.

| Figure | Sample | Objective lens | Illumination | Detector |
| --- | --- | --- | --- | --- |
| Fig. 2 | Silver mirror | 40x 1.15 NA (WI) | 660 nm LED | sCMOS camera |
| Fig. 3 | Silver mirror | 40x 1.15 NA (WI) | 660 nm LED | sCMOS camera,<br>PMT array |
| Fig. 4 | Live neurons | 25x 1.05 (SI) | 630 nm LED | sCMOS and<br>EMCCD camera |
| Fig. 5 | Live neurons | 40x 1.15 NA (WI) | 630 nm LED | sCMOS and<br>EMCCD camera |
| Extended Data Fig. 2b | Fluorescent<br>solution | 60x 1.49 NA (OI) | 630 nm LED | sCMOS camera |
| Extended Data Fig. 2d | Fluorescent<br>solution | 4x 0.13 NA, PhL | 630 nm LED | sCMOS camera |
| Extended Data Fig. 4 | Fluorescent<br>solution | 60x 1.49 NA (OI) | 630 nm LED | sCMOS camera |
| Extended Data Fig. 5a | Silver mirror | 40x 1.15 NA (WI) | 630 nm LED | sCMOS camera |
| Extended Data Fig. 5b | USAF target | 10x 0.45 NA | White LED | sCMOS camera |
| Extended Data Fig. 7 | Fluorescent<br>beads | 40x 1.15 NA (WI) | 630 nm LED | sCMOS camera |
| Extended Data Fig. 8 | Live neurons | 40x 1.15 NA (WI) | 630 nm LED | sCMOS and<br>EMCCD camera |

## Supplementary Section 1. Resolution of optical segmentation

The diffraction-limited resolution for optical segmentation at SLM is determined by the NA at the SLM (NA_SLM_), which is determined by the NA of objective lens (NA_obj_) divided by the magnification factor at the SLM (M_SLM_). In our design, the M_SLM_ is the product of the magnification of an objective lens (M_obj_: 40x) and that of optical relay to SLM (M_relay_ = 1.87x).

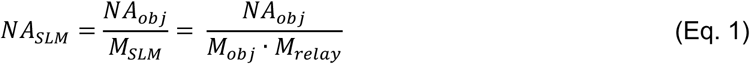

Applying the Equation 1, NA_SLM_ is ∼0.015 for 40x, 1.15NA objective lens and ∼0.012 for 63x, 1.4NA objective lens. By the Rayleigh criteria, the diffraction-limited resolution (d) is defined as

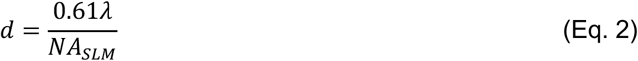

where λ is wavelength of emission (∼700 nm). Applying the Equation 2, the diffraction resolutions (d) are 28.5 μm and 35.6 μm for 40x and 63x objective lenses, respectively. Considering the rectangular-shaped field stop reducing the effective NA at the SLM (NA_SLM_) by ∼15%, the resolution (d) for 40x objective lens is ∼31 μm. The 3-level blazed grating pattern with the 9.2 μm pixel pitch of SLM provides an effective unit of optical segmentation of 27.6 μm, providing ∼55 μm of sampling-limited resolution at the SLM. This value corresponds to 735 nm for 40x objective lens, and 467 nm for 63x objective lens. Overall, the resolution for optical segmentation in this study is submicron-scale limited by the pixel pitch of SLM.

## Supplementary Section 2. Linear unmixing

To unmix inter-channel crosstalk caused by multiple diffraction orders, the diffraction efficiency matrix, *η*, is experimentally obtained. The relationship between measured intensity (î) and the efficiency matrix (*η*) can be expressed as

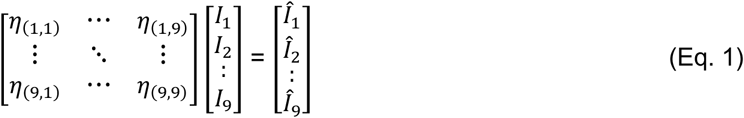

where *η*(i,j) is efficiency of diffraction from the ‘j’th channel to the ‘i’th channel, I_i_ is original intensity at the ‘i’th channel, and î_i_ is the measured intensity at the ‘i’th channel. We acquired intensity for each channel, î_i_, with uniform blazed grating patterns for 8-directions corresponding to the case when only I_i_ is nonzero or with no grating (i.e., only the I_5_ is nonzero). From the data, we measured η_(i,j)_ values. As expected, η_(i,j)_ value was negligible except for the +1st order diffraction (η_(i,j)_, where ‘i = j’), the 0th order diffraction (η_(i,j)_, where ‘i = 5’) and the −1st order diffraction (η_(i,j),_ where ‘i+j = 10’). By solving the linear equations between the paired channels (Eq. 2), we obtained I_i_ and used it for further analysis.

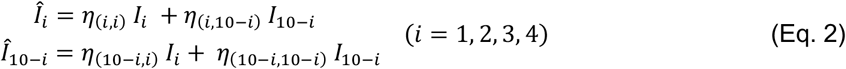

The residual 0th order signal is minimized by fine-tuning the diffraction efficiency matrix (η) based on the stepwise loss minimization (Eq. 3).

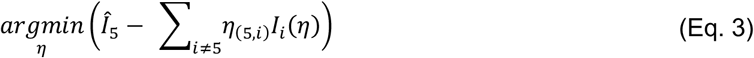

## References

1. Debanne, D., Bialowas, A. & Rama, S. What are the mechanisms for analogue and digital signalling in the brain? Nat. Rev. Neurosci. 14, 63–69 (2013).

2. Stam, C. J. & Straaten, E. C. W. van. The organization of physiological brain networks. Clin. Neurophysiol. 123, 1067–1087 (2012).

3. Fornito, A., Zalesky, A. & Breakspear, M. The connectomics of brain disorders. Nat. Rev. Neurosci. 16, 159–172 (2015).

4. Bassett, D. S. & Sporns, O. Network neuroscience. Nat. Neurosci. 20, 353–364 (2017).

5. Sasaki, T., Matsuki, N. & Ikegaya, Y. Action-Potential Modulation During Axonal Conduction. Science 331, 599–601 (2011).

6. Jayant, K. et al. Targeted intracellular voltage recordings from dendritic spines using quantum-dot-coated nanopipettes. Nat. Nanotechnol. 12, 335–342 (2017).

7. Tian, H. et al. Video-based pooled screening yields improved far-red genetically encoded voltage indicators. Nat. Methods 20, 1082–1094 (2023).

8. Abdelfattah, A. S. et al. Bright and photostable chemigenetic indicators for extended in vivo voltage imaging. Science 365, 699–704 (2019).

9. Piatkevich, K. D. et al. Population imaging of neural activity in awake behaving mice. Nature 574, 413–417 (2019).

10. Villette, V. et al. Ultrafast Two-Photon Imaging of a High-Gain Voltage Indicator in Awake Behaving Mice. Cell 179, 1590–1608 (2019).

11. Adam, Y. et al. Voltage imaging and optogenetics reveal behaviour-dependent changes in hippocampal dynamics. Nature 569, 413–417 (2019).

12. Hochbaum, D. R. et al. All-optical electrophysiology in mammalian neurons using engineered microbial rhodopsins. Nat. Methods 11, 825–833 (2014).

13. Platisa, J. et al. High-speed low-light in vivo two-photon voltage imaging of large neuronal populations. Nat. Methods 20, 1095–1103 (2023).

14. Liu, Z. et al. Sustained deep-tissue voltage recording using a fast indicator evolved for two-photon microscopy. Cell 185, 3408–3425 (2022).

15. Kannan, M. et al. Dual-polarity voltage imaging of the concurrent dynamics of multiple neuron types. Science 378, eabm8797 (2022).

16. Xiao, S. et al. Large-scale voltage imaging in behaving mice using targeted illumination. iScience 24, 103263 (2021).

17. Hoppa, M. B., Gouzer, G., Armbruster, M. & Ryan, T. A. Control and Plasticity of the Presynaptic Action Potential Waveform at Small CNS Nerve Terminals. Neuron 84, 778–789 (2014).

18. Cohen, C. C. H. et al. Saltatory Conduction along Myelinated Axons Involves a Periaxonal Nanocircuit. Cell 180, 1–12 (2020).

19. Wu, J. et al. Kilohertz two-photon fluorescence microscopy imaging of neural activity in vivo. Nat. Methods 17, 287–290 (2020).

20. Weber, T. D., Moya, M. V., Kılıç, K., Mertz, J. & Economo, M. N. High-speed multiplane confocal microscopy for voltage imaging in densely labeled neuronal populations. Nat. Neurosci. 26, 1642–1650 (2023).

21. Xiao, S., Giblin, J. T., Boas, D. A. & Mertz, J. High-throughput deep tissue two-photon microscopy at kilohertz frame rates. Optica 10, 763 (2023).

22. Wu, J., Ji, N. & Tsia, K. K. Speed scaling in multiphoton fluorescence microscopy. Nat. Photonics 15, 800–812 (2021).

23. Reynolds, S. et al. ABLE: An Activity-Based Level Set Segmentation Algorithm for Two-Photon Calcium Imaging Data. eNeuro 4, ENEURO.0012-17.2017 (2017).

24. Brondi, M. et al. High-Accuracy Detection of Neuronal Ensemble Activity in Two-Photon Functional Microscopy Using Smart Line Scanning. Cell Reports 30, 2567–2580.e6 (2020).

25. Zhang, Y. et al. Rapid detection of neurons in widefield calcium imaging datasets after training with synthetic data. Nat Methods 1–8 (2023) doi:10.1038/s41592-023-01838-7.

26. Giovannucci, A. et al. CaImAn an open source tool for scalable calcium imaging data analysis. Elife 8, e38173 (2019).

27. Cai, C. et al. VolPy: Automated and scalable analysis pipelines for voltage imaging datasets. Plos Comput Biol 17, e1008806 (2021).

28. Grinvald, A., Ross, W. N. & Farber, I. Simultaneous optical measurements of electrical activity from multiple sites on processes of cultured neurons. Proc. Natl. Acad. Sci. 78, 3245–3249 (1981).

29. Wang, T. et al. Image sensing with multilayer nonlinear optical neural networks. Nat. Photonics 17, 408–415 (2023).

30. Li, X. et al. Real-time denoising enables high-sensitivity fluorescence time-lapse imaging beyond the shot-noise limit. Nat. Biotechnol. 41, 282–292 (2023).

31. Li, X. et al. Reinforcing neuron extraction and spike inference in calcium imaging using deep selfsupervised denoising. Nat. Methods 18, 1395–1400 (2021).

32. Lecoq, J. et al. Removing independent noise in systems neuroscience data using DeepInterpolation. Nat. Methods 18, 1401–1408 (2021).

33. Eom, M. et al. Statistically unbiased prediction enables accurate denoising of voltage imaging data. Nat. Methods 20, 1581–1592 (2023).

34. Zhou, P. et al. Efficient and accurate extraction of in vivo calcium signals from microendoscopic video data. Elife 7, e28728 (2018).

35. Xie, M. E. et al. High-fidelity estimates of spikes and subthreshold waveforms from 1-photon voltage imaging in vivo. Cell Reports 35, 108954 (2021).

36. Gustafsson, M. G. L. Nonlinear structured-illumination microscopy: Wide-field fluorescence imaging with theoretically unlimited resolution. Proc. Natl. Acad. Sci. 102, 13081–13086 (2005).

37. Dan, D. et al. DMD-based LED-illumination Super-resolution and optical sectioning microscopy. Sci .Rep. 3, 1116 (2013).

38. Emiliani, V., Cohen, A. E., Deisseroth, K. & Häusser, M. All-Optical Interrogation of Neural Circuits. J. Neurosci. 35, 13917–13926 (2015).

39. Bounds, H. A. et al. All-optical recreation of naturalistic neural activity with a multifunctional transgenic reporter mouse. Cell Rep. 42, 112909 (2023).

40. Adesnik, H. & Abdeladim, L. Probing neural codes with two-photon holographic optogenetics. Nat. Neurosci. 24, 1356–1366 (2021).

41. Papagiakoumou, E. et al. Scanless two-photon excitation of channelrhodopsin-2. Nat. Methods 7, 848–854 (2010).

42. Papagiakoumou, E., Ronzitti, E. & Emiliani, V. Scanless two-photon excitation with temporal focusing. Nat. Methods 17, 571–581 (2020).

43. Shemesh, O. A. et al. Temporally precise single-cell-resolution optogenetics. Nat. Neurosci. 20, 1796–1806 (2017).

44. Bowman, A. J., Huang, C., Schnitzer, M. J. & Kasevich, M. A. Wide-field fluorescence lifetime imaging of neuron spiking and subthreshold activity in vivo. Science 380, 1270–1275 (2023).

45. Brinks, D., Klein, A. J. & Cohen, A. E. Two-Photon Lifetime Imaging of Voltage Indicating Proteins as a Probe of Absolute Membrane Voltage. Biophys. J. 109, 914–921 (2015).

46. Sankaran, J. et al. Simultaneous spatiotemporal super-resolution and multi-parametric fluorescence microscopy. Nat. Commun. 12, 1748 (2021).

47. Singh, A. P. et al. The performance of 2D array detectors for light sheet based fluorescence correlation spectroscopy. Opt. Express 21, 8652–8668 (2013).

48. Walker, A. S. et al. Optical Spike Detection and Connectivity Analysis With a Far-Red Voltage-Sensitive Fluorophore Reveals Changes to Network Connectivity in Development and Disease. Front. Neurosci. 15, 643859 (2021).

